# Artificial neural networks enable genome-scale simulations of intracellular signaling

**DOI:** 10.1101/2021.09.24.461703

**Authors:** Avlant Nilsson, Joshua M. Peters, Bryan Bryson, Douglas A. Lauffenburger

## Abstract

Mammalian cells adapt their functional state in response to external signals in form of ligands that bind receptors on the cell-surface. Mechanistically, this involves signal-processing through a complex network of molecular interactions that govern transcription factor (TF) activity patterns. Computer simulations of the information flow through this network could help predict cellular responses in health and disease. Here we develop a recurrent neural network constrained by prior knowledge of the signaling network with ligand concentrations as input, TF activity as output and signaling molecules as hidden nodes. Simulations are assumed to reach steady state, and we regularize the parameters to enforce this. Using synthetic data, we train models that generalize to unseen data and predict the effects of gene knockouts. We also fit models to a small experimental data set from literature and confirm the predictions using cross validation. This demonstrates the feasibility of simulating intracellular signaling at the genome-scale.

## Introduction

The healthy body continuously adapts to the environment by altering the molecular state of its cells. This primarily occurs through binding of multiple types of ligands to receptors on the cell-surface, this acts as signals that are propagated through molecular interactions culminating in activation of transcription factors (TF) and subsequent transcription of genes. Rather than constituting independent paths from receptors to specific genes, signaling is conducted through a complex network with spatial and temporal components^1^. This enables the cell to compute a response to stimulation with multiple ligands^2,3^, e.g. co-stimulation of human macrophages gives rise to a spectrum of cellular activation states^4^. Disruptions to the network can cause disease, e.g. activating mutations in the signaling protein BRAF is present in 40- 50% of all melanoma tumors^5^, i.e. skin cancer, and single target treatments are not always sufficient due to cellular adaptations, e.g. tumors often acquire resistance to BRAF-inhibitors^5^. A systems perspective on signaling is required to better understand responses to co-stimulation and predict the effects of drugs. Such an understanding could be obtained through genome-scale computer simulations of signaling that have long been anticipated^6–8^.

By now, many requisites for genome-scale models of signaling are in place. The network topology has been extensively characterized with thousands of biochemical interactions collected in databases^9^ and with visual maps available for many signaling pathways, e.g. through the Kyoto Encyclopedia of Genes and Genomes (KEGG)^10^. Genome wide data can be generated using high-throughput methods, e.g. activities of hundreds of TFs can be statistically inferred from transcriptomics data^11^ and cellular responses to combinations of ligands, can be characterized through co-stimulation experiments^2^. For metabolism genome-scale simulations are routinely performed using the flux balance analysis (FBA) framework, which predicts intracellular fluxes using steady state assumptions, linear optimization and data on metabolic exchange rates^12^. It has been used to gain system level insight on a wide range of topics, e.g. the effect of intercellular compartmentalization on the flux of glutamate in cancer^13^ or the influence of metabolic trade-offs on oxygen consumption in muscle cells^14^. However, the linear FBA methodology cannot be applied to signaling, in which nonlinear relationships are typically important to capture and stoichiometric constraints are less straightforward to impose.

Current signaling models are often based on ordinary differential equations (ODE) or logic rules^7,8,15^ and face challenges when expanding to the genome-scale^7,16^. Yet, several of these have been overcome by simplifying assumptions. Explicit enumeration of microstates, which has been successful for individual proteins, is numerically intractable at the genome-scale^17^ due to a combinatorial explosion of states from posttranslational modifications and protein complexes. This is circumvented by models that omit enumeration, e.g. signal flow models represent signaling as a signed directed graph with scalar activity values for each signaling molecule^15^. Cellular activity occurs across multiple timescales, e.g. conformational changes of proteins occurs much faster than signaling events, while protein translation from mRNA occurs much slower. The requirement by network-wide models for simulation of long time-courses at high resolution can be overcome using quasi-steady-state approximations^17,18^ that assume that faster processes are instantaneous and slower processes as constant. However, two major limitations remain for reaching the genome-scale using current methods: predefined equations are needed for each molecule, while the exact mechanism is often unknown; and parameter estimation may require problematically long computational times for the largest models despite major advances^19^. Therefore, an alternative framework for modeling signaling may be warranted.

Advancements in artificial neural networks (ANN) have enabled large-scale models in many different areas, including drug discovery and genomics^20^. ANNs approximate unknown and highly complex functions through a sequence of linear matrix operations and non-linear transformations. These approximations, sometimes containing millions of parameters, can be rapidly trained from paired samples of input and output data using the backpropagation algorithm^20,21^. While ANNs excel at predictions, their underlying mechanism is often elusive and therefore more interpretable ANNs based on prior knowledge have been proposed for modeling biological systems^22^. For example, a feed forward neural network (FFNN) with a network topology derived from known signaling interactions has been used to predict cell types from gene expression data^22^. However, FFNN do not allow feedback loops, which are frequent in signaling, and therefore recurrent neural networks (RNN) may be a more suitable architecture for modeling signaling networks. It has previously been shown that a an RNN without prior knowledge constraints can recapitulate the output of a small ODE-model of signaling^23^.

Here we construct a framework for rapid parameterization and simulation of intracellular signaling using RNNs. We first construct an activation function suitable for approximating the steady state behavior of different molecular mechanisms. We then introduce a sparse RNN formalism that encodes the topology of a known signaling network. The RNN uses ligand concentrations as input to predict steady state TF activities and we construct a regularization function that ensures that steady state is reached. To test the data requirements for training generalizable models, we generate synthetic data from a reference model with computationally derived parameters. Models trained on modestly sized (400-800 samples) synthetic datasets, accurately predict most randomly generated input-output pairs from the reference model. Additionally, the trained model predicts the effect of simulated gene knock outs (KO). To demonstrate the frameworks applicability to real world data, we fit a model using a small transcriptomics dataset from literature involving macrophages stimulated with different combinations of ligands. We discuss how genome-scale signaling models may leverage new types of high throughput data and facilitate personalized medicine.

## Results

### Approximating molecular interactions at steady state

For the purpose of the signaling framework developed herein, molecular interactions are assumed to always be at steady state. This can be justified by timescale separation, as these events are expected to occur on the order of milliseconds compared to signal transduction that evolves over several minutes. Molecular dynamics here signifies interactions between signaling molecules through a range of different mechanisms, e.g. phosphorylation, binding, or conformational changes. The steady state assumption implies that the activity of the target molecule of the interaction is a single valued function of its source molecules that are considered constant at that instant. This activity depends on the specific molecular mechanism (Fig. 1a) with the simplest arguably being independent activation and inhibition that may be interpreted as phosphorylases and phosphatases respectively.

**Figure 1.**
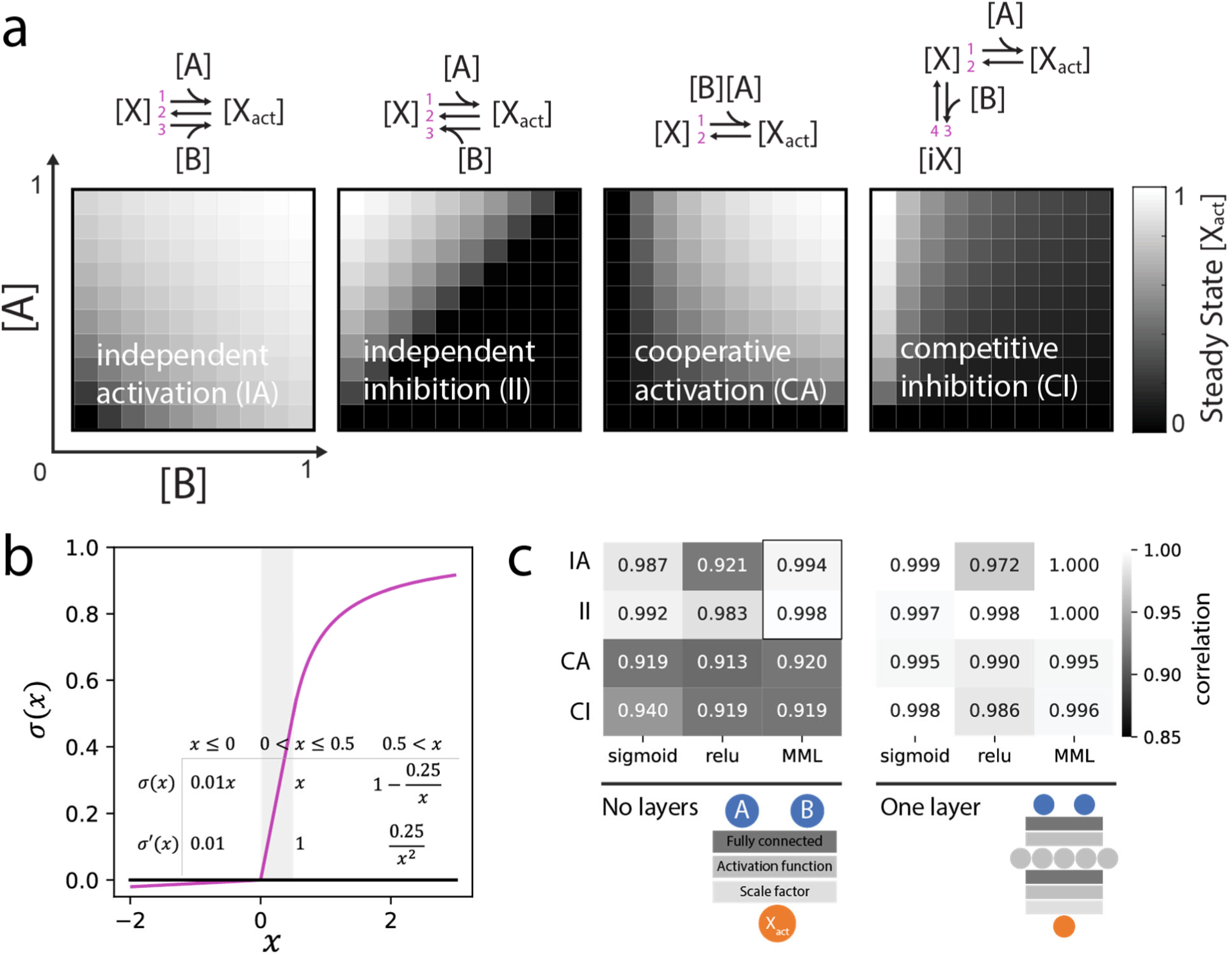
Modeling molecular mechanisms using a FFNN. **a)** The steady state for different mechanisms involving two source molecules (A and B) at constant concentration and a target molecule with an inactive state (X) and active state (X_act_). Results were attained by running ODE simulations until steady state, reaction rates (1-4) were parameterized manually (values in Supplemental Table S1), maximum activity was scaled to 1. **b)** The Michaelis- Menten-like (MML) activation function is designed as a monotonously increasing, continuous function with a maximum of 1 and continuous first derivative (except at 0). It is composed of three segments; a leaky; a linear; and a saturating. The leaky and linear segments correspond to the leaky ReLU activation function, as the Michaelis Menten equation is not defined for negative input. The saturating segment is composed of a shifted and scaled Michaelis-Menten equation. **c)** FFNNs models of molecular mechanisms with different activation functions and number of layers. Model were trained on steady state activity for a grid of 7×7 linearly spaced source concentrations and tested on a 20×20 grid, the Pearson correlation between prediction and test data was calculated (mean of three runs). A black outline marks the performance of MML for independent activation and inhibition with no hidden layers.

In many cases the exact molecular mechanism of a signaling interaction will be unknown, but its input-output relation can be approximated by a neural network. Directed acyclic graphs, i.e. a FFNN, are appropriate models for interactions that are assumed to instantaneously reach steady state^17^ and for independent activation and inhibition there is a direct mapping between their analytical steady state solution and a FFNN with the Michaelis--Menten equation as activation function (Supplementary Fig. S1a). Based on this we developed a problem-specific activation function, the Michalis-Menten like (MML) activation function (Fig. 1b) with two main features; preventing negative states that would be non-physiological; and preventing states >1 that are non-physiological assuming that this represents full saturation. Physiological constraints are thus imposed at the level of the activation function, allowing weights and biases to take on arbitrary values. In practice the MML was taken as the leaky version of the Rectified Linear Unit (ReLU) activation in its standard formulation^24^ for negative inputs. This prevents a strict 0 gradient that may cause irrecoverable inactivation of nodes during training leading to blocked signaling in sparse networks. The MML was taken as ReLU also for input values less than 0.5 to allow a range where signaling states can be passed forward without alteration.

We found that a FFNN with this activation function and no hidden layers provided a good approximation of independent activation/inhibition (Fig. 1c) outperforming the other activation functions that were tested. The overall performance was acceptable also for other molecular mechanisms, although prediction errors were not uniformly distributed (Supplementary Fig. S1b). An advantage of the MML model without hidden layers was that the sign of weights directly corresponded to the mode of action (MOA), activation (positive) or inhibition (negative). This allows for a straight forward implementation of MOA-constraints. Additionally, it requires markedly less calculations than multilayered FFNNs. For FFNNs with one hidden layer all of the tested activation functions produced excellent approximations (Fig. 1c).

### Constraining a recurrent neural network with prior knowledge of signaling

Signaling involves a network of molecular interactions whose effects propagates over cellular distances from receptors at the surface to TFs in the nucleus. In order to represent these interactions, which include feedback loops, a sparse RNN formulation was developed as a model of cellular signaling. We constructed a minimal signaling network to demonstrate the framework (Fig. 2a). The structure of this prior knowledge network was encoded by a sparse matrix holding the weights of its molecular interactions (Fig. 2b). The overall expression, also known as a first order non-linear difference equation, iteratively calculates the signaling state from the state at the previous timestep and includes ligand concentrations as input and a bias term, which may be interpreted as basal activation or thresholding. In this study molecular interactions were modeled without hidden layers so that MOA could easily be constrained, but the approximations of molecular interactions could have been made arbitrarily complex by adding intermediary nodes between sources and targets.

**Figure 2.**
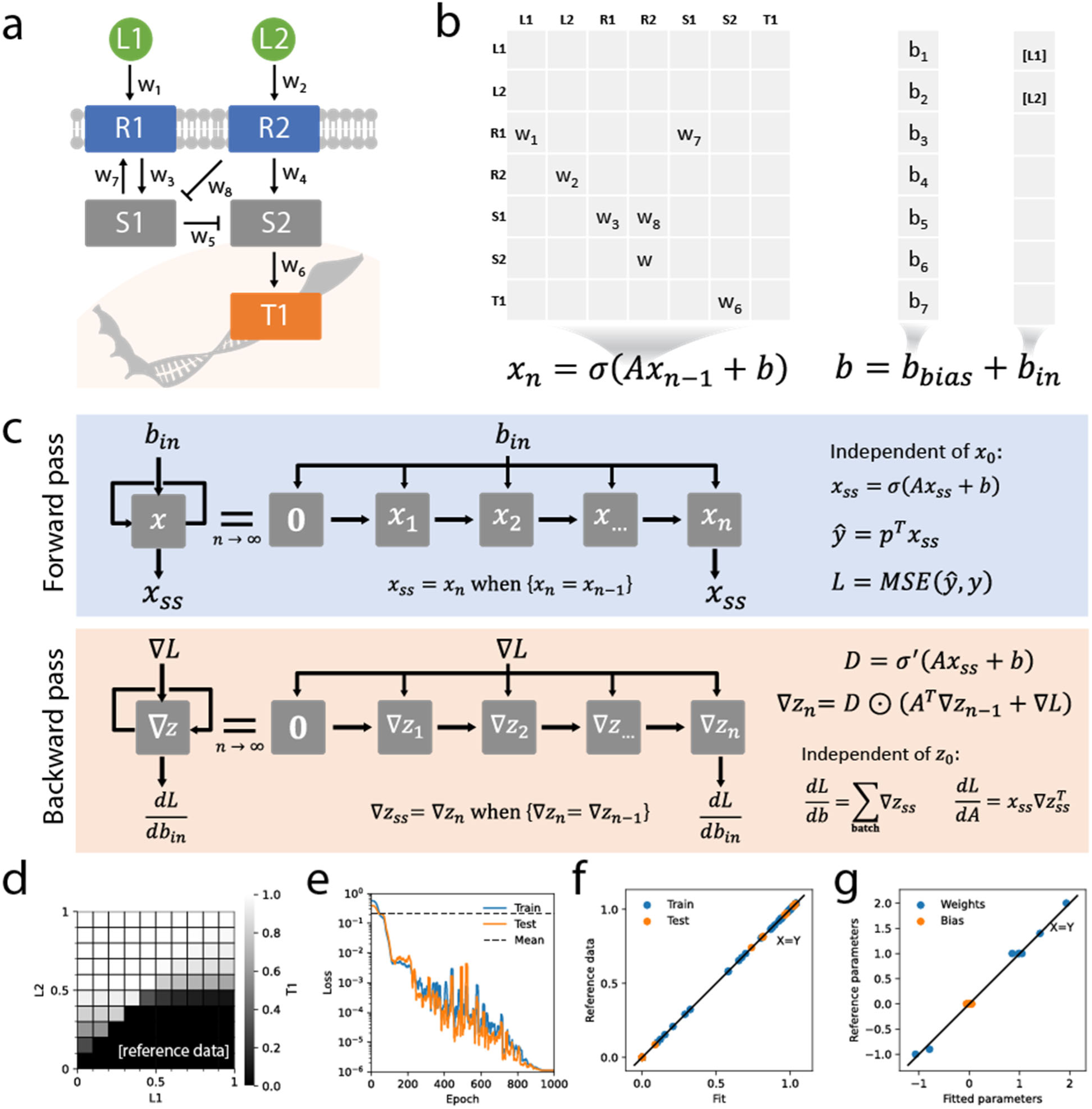
Modeling a simple signaling network using an RNN. **a)** A simple signaling network including ligands (L1, L2) receptors (R1, R2), signaling proteins (S1, S2) and a TF (T1) interconnected by 8 interactions (w_1_, …, w_8_). **b)** The difference equation calculates the state (*x*_*n*_) at timestep n from the previous time step. The matrix A holds the interaction weights and the bias term (*b*_*bias*_) and input term (*b*_*in*_) are added, the activation function (*σ*) is applied at each timestep. **c)** Calculations are repeated until a steady state (*x*_*ss*_) is reached and the predicted TF activity (*ŷ*) is projected (p) from the steady state. Loss (L) is calculated by comparison to reference data (y), e.g. as mean square error (MSE). It is back propagated to provide the partial derivatives with respect to weights and biases at steady state. Here ⊙ is element wise multiplication. To prevent potential exploding gradients, the loss is clipped at each step (see methods). **d)** Reference data generated by a parameterized model. **e)** A model trained on the reference data by minimizing the loss using stochastic gradient decent. Loss from predicting the mean value for comparison (dashed line). **f)** Perfect fit (Train) and generalization (Test) to reference data. **g)** Parameters in in agreement between fit and reference.

It is here assumed that signaling activity reaches steady state after evolving for some predefined number of timesteps (Fig. 2c) and TF-activities are projected from the steady state. Ligand-concentrations, weights, and biases are all assumed to stay constant during the iterations. This can be motivated by time- scale separation, although regulatory events, which take place over hours, are expected to result in translation of proteins that are excreted as ligands or remodel the network interactions. Often RNNs are used to fit time series or other sequence data, but here intermediate states are discarded resulting in a one-to-one relation between ligand patterns and steady state TF activities. It can be noted that while internally, a trajectory is computed from some initial state (here all zeros) to steady state, the steady state does not explicitly depend on the initial state or any of the intermediary steps and these are therefore not required to reflect biologically relevant transitions. Two other implications of the steady state assumption are; that any oscillations exhibited by the network are dampened; and that switch-like behavior, where network responses depend on the initial state, must be encoded as differences in weights or biases.

When training a model using this framework, any potential prediction errors can be back propagated to adjust the model parameters. The unrolling of an RNN into discrete timesteps is commonly referred to as backpropagation through time (BPTT)^25^. Here the BPTT expression is simplified by the steady state assumption and the assumption of constant input (Fig. 2c). Due to these assumptions the gradients only depend on the steady state values and due to vanishing of gradients from early timesteps, the back- propagated error can be assumed to reach a steady state that is independent from the trajectory by which it is computed (See Supplementary Fig. 2 for a numerical comparison and Supplementary methods for derivation). It can be noted that BPTT, for this restricted RNN, strongly resembles loopy belief propagation that is used for Bayesian inference on cyclic graphs^26^, where error messages are propagated until convergence.

Models can be constructed on training data, then tested for generalization on previously unseen data. The data, containing ligand–TF activity pairs, was generated from a reference model (Fig. 2d) with manually assigned parameters. A model was trained using this data, i.e. without direct access to the parameter values. Terms were added to the loss function to constrain weights by their MOA and constrain biases at ligand positions to zero, since their concentration was assumed to be provided as input (see methods for implementation). Additionally, regularization terms controlling the L2 norm of the parameters were added to prevent overfitting, as is common practice. The model was trained using the ADAM optimizer^27^ with a cosine learning rate schedule and warm restarts, as has been proposed by others^28^. Using this setup, it was possible to train (Fig. 2e) a model to a near perfect accuracy, both on data used for training (80%), and on test data (20%) that was left out of the training set at random (Fig. 2f). We tested the methods sensitivity to non-uniformly distributed training data by adversely selecting samples that were left out of training (supplementary Fig. 3a) and this reduced generalization marginally, e.g. removing the bottom left quadrant reduced the correlation (Pearson) of predictions to 0.85.

For this particular model structure, the trained model accurately recovered the original parameter values (Fig. 2g). However, it was possible to construct a network, with sequential nodes without branching, where this did not occur (Supplementary Fig. S3b) even though the network generalized perfectly to test data, i.e. the parameters were not identifiable. Nevertheless, there was a strong correlation between the predicted state vector of trained- and reference-model, suggesting that the learned model may be able to accurately predict the effects of perturbating states, despite the inconsistent parametrization.

### Constraining the spectral radius to enforce steady state

Feedback loops can prevent an RNN from reaching steady state. The formulation above assumes that a steady state is reached within a specified number of timesteps. However, depending on the parametrization this may not occur. To not reach steady state, could yield non-sensical output and may also be detrimental to gradient calculations, preventing training convergence. The requirement to reach steady state can be expressed formally using eigenvalue analysis of the linearized difference equation (Fig. 3a). For the model to eventually reach steady state, the absolute value of the largest eigenvalue of the transition matrix, i.e. the spectral radius, must be less than 1 (See supplementary methods for derivation). Similar ideas have previously been explored for linear systems^29^ and for RNNs^30,31^.

**Figure 3.**
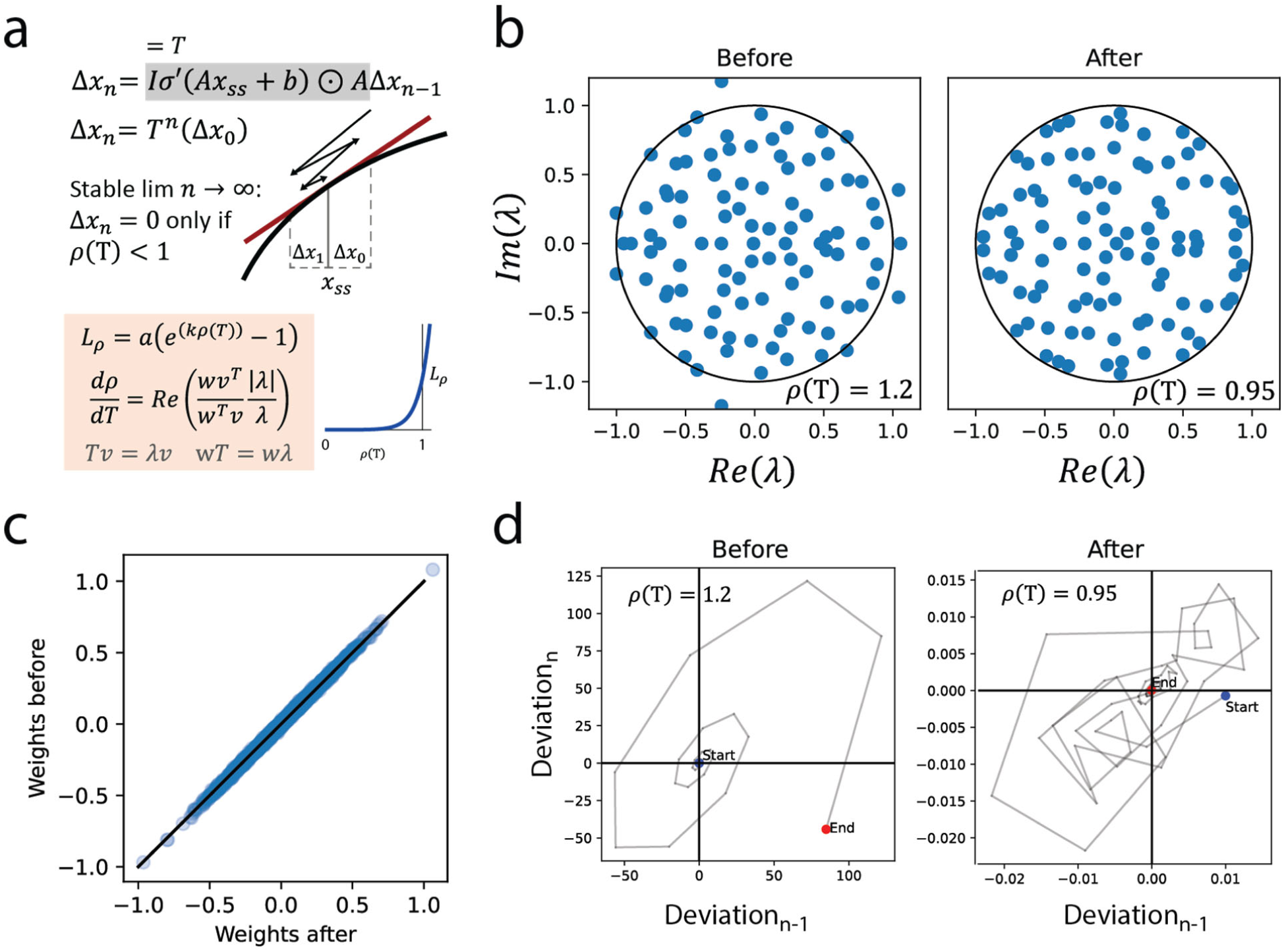
Stability of each steady state is controlled by the spectral radius of a transition matrix. **a)** Linearization of the difference equation (first order Taylor expansion) around a steady state (*x*_*ss*_) and shift of coordinates with the steady state as origin yields a homogeneous linear difference equation with transition matrix (T), here *l* is the identity matrix. Stability requires that a deviation (Δ*x*_0_) from steady state tends to 0 for repeated multiplication by T. This occurs only if the spectral radius (*ρ*) of T, i.e. the eigenvalue (*λ*) of T with largest absolute value, is less than 1. A regularization term (*l*_*ρ*_) is constructed to constrain the spectral radius. Its gradient is a function of the eigen value and left (*w*) and right (*v*) eigenvectors. Note that the imaginary component of this function is orthogonal to the radius and can be ignored. The Re function elementwise returns the real part of complex numbers. **b)** All eigenvalues of a 100×100 transition matrix before and after a reduction of the spectral radius with the loss function using gradient decent for 200 steps. Matrix parametrized at random with 20% non-zeros. **c)** Strong similarity between matrix weights before and after reduction suggesting that the regularization will cause minimal disruption to the learning. Trajectories are stable after shrinking the transition matrix, here the trajectory of the first element.

It is possible to constrain the spectral radius. Its partial derivatives can be computed (for a numerical demonstration see supplementary Fig. S4) since it is a locally smooth function of the weights^32^. We introduced a regularization term to control the spectral radius using gradient decent (Fig. 3b) and with marginal effects on the magnitudes of the weights (Fig. 3c) it ensures steady state behavior (Fig. 3d). The introduction of the spectral radius in the loss function can be viewed as imposing a prior on the temporal complexity of the model. It should be noted that while the spectral radius regularization ensures that all conditions in the training data reach steady state, it does not guarantee that this holds for arbitrary conditions, i.e. untested conditions may be unstable. With this, we had the prerequisites to simulate networks of arbitrary size and wiring.

### Parameterizing a large model for synthetic data generation

To put the framework to the test, we reconstructed a more comprehensive signaling network. For this, we turned to an online database, OmniPath^9^, that collects evidence of signaling interactions in human cells. The full set of interactions in OmniPath is very comprehensive and includes both well-characterized interactions and results from single high-throughput experiments. To ensure a model of high-quality, we used a subset of interactions that listed KEGG^10^ as reference database and for which the MOA was a known (Fig. 4a). Nodes were labeled as ligands, receptors, signaling molecules or TFs, based on annotation in OmniPath (see methods).

**Figure 4.**
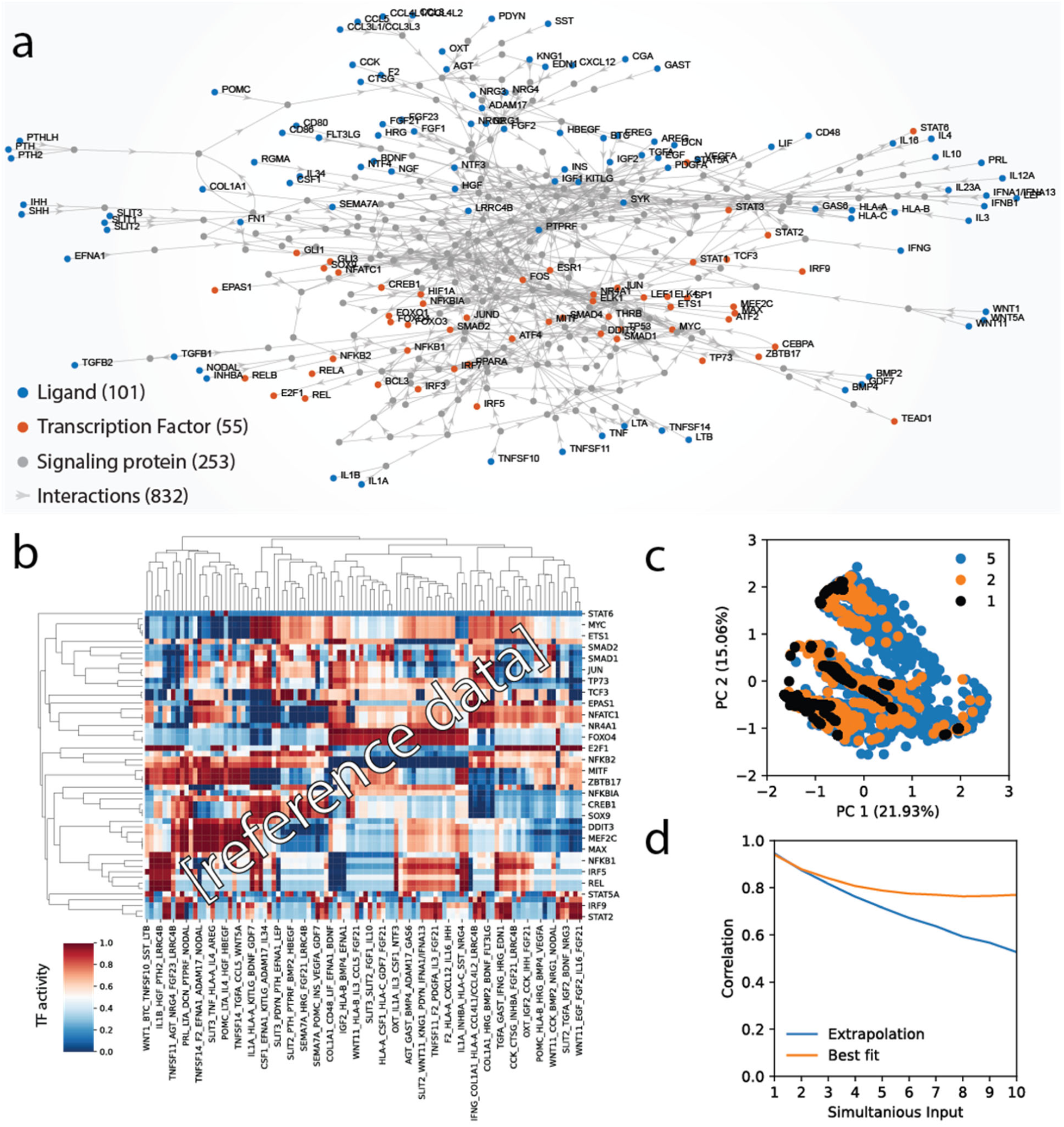
Synthetic data from automatically parameterized model. **a)** A large signaling model was reconstructed and automatically parameterized by maximizing an objective function. **b)** Synthetic data (N=100) generated from the model for combinations of 5 different ligands. **c)** Increasing combinations of ligands increases the area of TF-patterns in principal component space, 2000 randomly sampled conditions per combination-level. **d)** TF activity is not a linear combination of the activity of individual ligands. Extrapolation of a linear model trained on single ligands and best fit linear models trained on data from multiple ligands (20-fold cross validated).

We set up a reference model to generate synthetic data. To parameterize such a large model by hand would be daunting, in particular as the output of a meaningful model should involve complex integration of the input. To overcome this, we devised a setup to automatically generate parameters based on desired properties of the model (see methods). Briefly, using randomly generated input, an objective function was optimized to simultaneously minimize; mean correlation between conditions and TFs; the L2 norm of biases and weights; and deviations from a uniform distribution of activities for TFs and conditions. Additionally, the spectral radius, MOA of weights and the bias term on ligands were constrained. The resulting model generated rich synthetic data with biologically plausible TF activity patterns (Fig. 4b). Principal component analysis of the models TF-patterns showed that increasing the number of ligands increased the covered space (Fig. 4c), consistent with complex interactions and emergent states. This was also supported by generalized linear modeling of the patterns (Fig. 4d) that showed a decreasing fit for increasing number of simultaneous inputs. This parameterized reference model demonstrates the computational capacity that lays latent in the topology of the signaling network.

The time complexity of the framework affects its practical feasibility to learn large networks. Network size can be characterized by the number of signaling nodes (n) or by the number of non-zero interactions (z) and the bottleneck involves sparse matrix multiplication between the weight matrix and the state vector with a naïve complexity of zn^2^, meaning that simulation time increases linearly with the number of interactions but that doubling the number of nodes requires 4 times longer simulation time. For biologically relevant networks with between 1000 and 19000 nodes and ∼10 interactions per node, we observe a linear increase in wall time from 0.02 to 0.2 seconds per pass (Supplementary Fig. 5). However, the purpose of the algorithm is to train generalizable models from data, and the amount of data and number of epochs of training required may also depend on the network size (see supplementary methods for a more in-depth analysis). So, while the complexity is well defined for simulating a condition, i.e. a forward pass, the complexity of training a generalizable model, so far remains an empirical question, although polynomial bounds on the number of epochs have been established for some classification tasks using RNNs^33^.

### Training generalizable models on synthetic data

To test the data requirements for generating generalizable models, we trained models on synthetic data generated from the reference model (Fig. 5a). To aid in the generalization and prevent the model from getting stuck in local minima during training several regularization techniques were applied (see methods). Briefly, the state variable was regularized to have approximately uniform distribution and a non-negative max value across conditions; weights were regularized to have non-zero values and L2 regularization was applied to all parameters; gaussian noise was added to the state variable with the level of noise decayed throughout the training in proportion to the learning rate. Training with noise could be considered a more biologically realistic alternative to drop-out, a regularization technique that aims to decreases the dependency on specific nodes by removing them at random. Experiments with drop-out on knowledge primed neural networks by others^22^ showed that a much lower dropout rate than the default (50%) is required, presumably due to the likelihood of complete blockage when the number of possible paths are limited.

**Figure 5.**
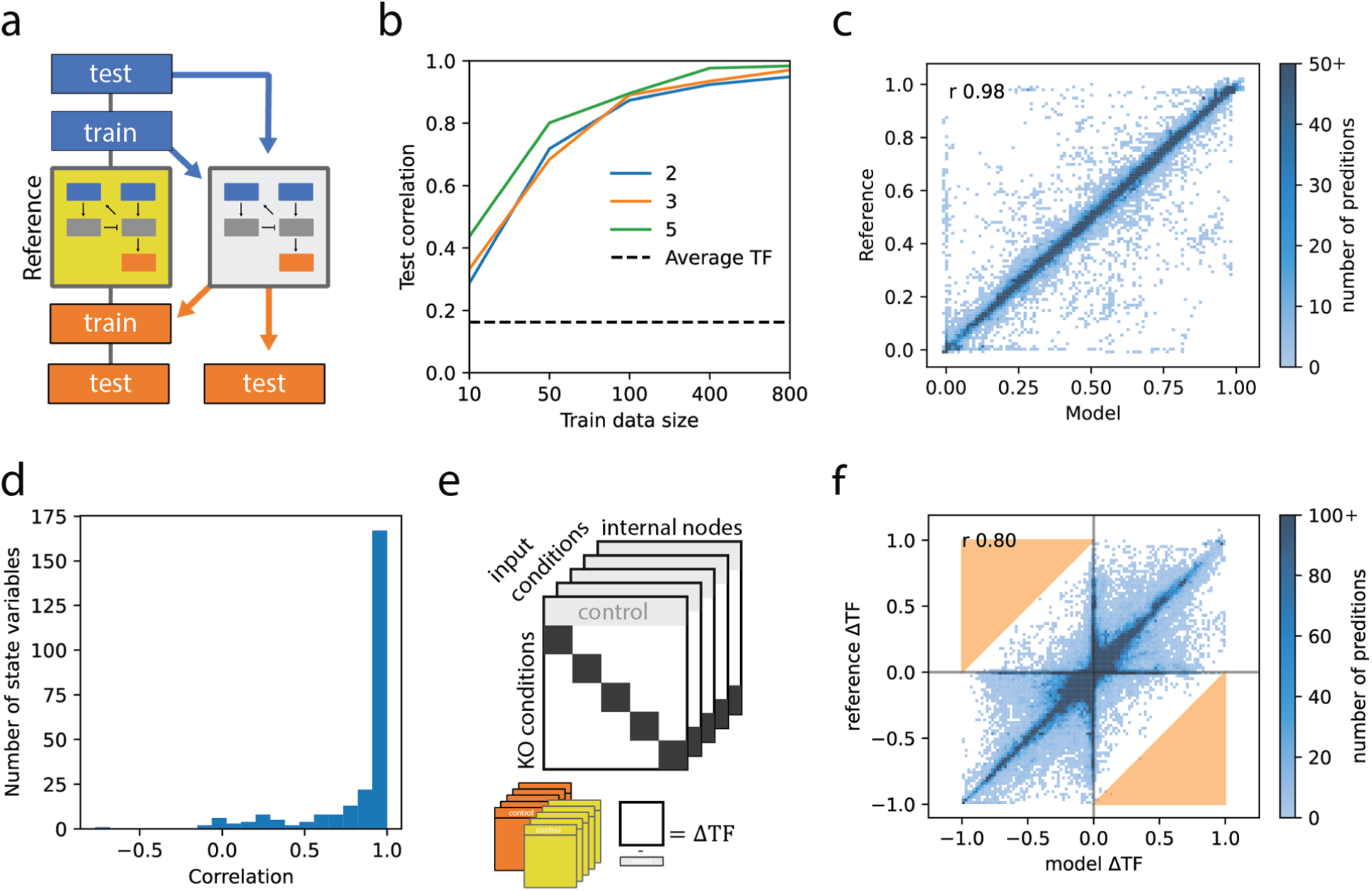
A model trained on modest amounts of synthetic data generalizes well. **a)** Synthetic training data was generated from a reference model and used to train an independent model. **b)** Training data with different numbers of conditions (10, 50, 100, 400, 800) and simultaneous ligands (2, 3, 5). Generalization performance was evaluated on 1000 independently generated test conditions as the Pearson correlation between prediction and reference. A non-zero correlation was attained for predicting the average of each TF. Training was conducted for 10000 epochs with a batch-size of 5. **c)** Comparison of model prediction and reference for the best performing model. Histogram truncated at 50 to emphasize exceptions. **d)** Mostly high correlation between states of model and reference, here only internal nodes, i.e. excluding ligands and TFs, were evaluated since they are not directly provided by the data. **e)** KO of a signaling node was simulated by applying a strong negative bias to the node, resulting in their state being close to zero after applying the activation function. **f)** Independent KO of 253 internal nodes under 100 random conditions. The change in TF activity for KO compared to no KO (control) was in good agreement between model and reference. For predictions to be off by more than 1 (marked in orange) both KO and control must be incorrectly predicted. Histogram truncated at 100.

With these techniques we fit models that generalized to a quite favorable extent. The amount of data required for this was investigated by training models with increasing amounts of randomly generated conditions for different numbers of simultaneous ligands (Fig. 5b). More simultaneous ligands improved generalization, and excellent performance was attained at the highest data settings. As expected, training models without spectral radius regularization caused training to diverge, resulting in poor fits (Supplemental Fig. S6a). A low, but non-zero, correlation was attained for models trained on data with scrambled order of conditions (Supplementary Fig. S6b). This could be due to the model learning general differences between distributions of individual TFs and was corroborated by an even higher correlation from taking the average of each TFs as prediction. We were concerned about potential information leakage from the reference model, since some of the regularization terms were shared with the parameterization algorithm, but training a model using only regularization terms (without fitting to data) did not perform better than predicting the average of each TF (Supplementary Fig. S5b), suggesting that leakage was not substantial.

For the best model the predicted TF values generally fell on the line of identity when comparing to reference (Fig. 5c). There were however some notable exceptions, these corresponded to a few poorly predicted conditions with correlations as low as 0.2 whereas the correlations of individual TFs were all above 0.9 (Supplementary Fig. S6c). Training with additional data could potentially alleviate this issue, since a larger state space would be sampled, but the saturating trend in generalization after 400 samples (Fig. 5a) suggests that perhaps further improvements to the regularization may be more economical. We found that, in general, parameters were not identical between reference and trained models (Supplementary Fig. S6d), presumably due to lack of identifiability, but that most of the state variables were still highly correlated between trained and reference models (Fig. 5d).

We hypothesized that the fitted model would predict *in silico* knock outs (KO) of signaling molecules in the reference model without training on such data. If successful, this would mean that the trained models had acquired the same structural dependencies as the reference model. For models trained on data from living cells, this would correspond to the ability to predict systemic effects of mutations or drugs. We simulated KO of each of the signaling molecules under several different conditions, i.e. in presence of different ligands. Although many KOs had limited impact on most TFs, the predicted difference in TF activity was similar between reference and fitted models (Fig. 5e), meaning that KO events were in general successfully predicted.

### Predicting signaling in ligand stimulated macrophages

In order to apply the framework to actual experimental cell biology data, TF activities must be estimated for each condition of cell stimulation or perturbation. For this we used a gene set enrichment based method, Dorothea^11^, that estimates probabilities of TF activities from mRNA concentrations of their target genes. Given that the lifetime of mRNA is expected to be much shorter than regulatory changes in transcription rates^34^, mRNA concentrations can be expected to be proportional to their formation rates and thus reflect the activity of the TFs that regulate their expression. A potential limitation with this approach is that the statically inferred probabilities of activation may not have a direct biological interpretation. Here they are taken as fractional activation that may affect the rate of transcription through time-occupancy at TF binding sites or by recruitment of polymerases, which may depend on multiple factors e.g. binding affinity, concentration, signaling state. It can be expected that there exists some relation between these activities and the probabilities inferred from differences in mRNA concentrations, however, this relation may be non-linear and noisy, which may partially be accounted for by the parametrization of the RNN model.

A transcriptomics dataset from ligand-stimulated macrophages was retrieved from literature^4^. Dorothea was applied to estimate probabilities of TF activation (Fig. 6a). There was in general good agreement among the biological replicates, but for four TFs the activities across replicates were deemed too noisy, and they were excluded from further analysis, notably STAT2 and IRF9 (Supplementary Fig. S7a). Some differences in variability were also observed among conditions (Supplementary Fig. S7b) but none were excluded. The inferred activity patterns appear to largely agree with known biology. For example, the transcription factors RelA and RelB are part of the NF-κB signaling cascade and induced by inflammatory ligands, e.g. interferons, lipopolysaccharide (LPS) and TNFα^35^. The ligands Il4 and IL13 display similar TF-activity profiles and are opposed to the inflammatory ligands, which is expected since they both signal through IL4R and are known to induce an anti-inflammatory (M2) response^36^. The presence of these ligands is here associated with SMAD3 activity, which may be a secondary effect from secreted TGFβ1^36^. The observed differences in TF-patterns between standard (LPSc) and ultra-pure (upLPS) LPS-qualities, are somewhat unexpected, but may potentially be explained by activation of TLR2 by impurities alongside the expected TLR4 activation^37^. Differences in signaling outcome for these two receptors have previously been noted^35^.

**Figure 6.**
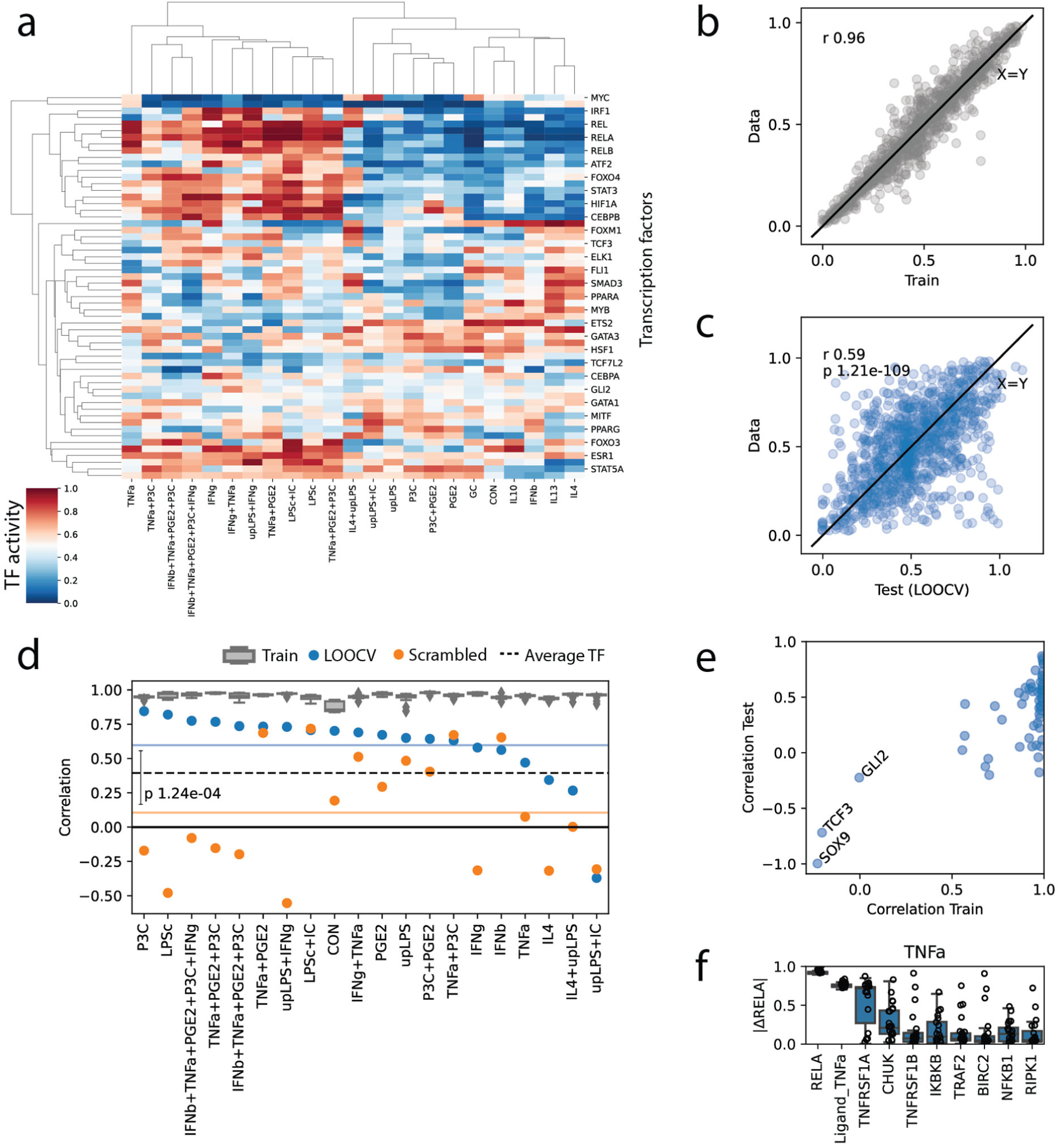
Model applied to experimental data from literature. **a)** An experimental dataset was retrieved from literature consisting of 23 conditions with combinations of 12 different ligands. Activity was estimated for 58 transcription factors and a model was reconstructed to fit the data consisting of 1069 signaling nodes and 5886 interactions. **b)** A good fit (r = Pearson correlation) was attained within 35 min wall time on a laptop. **c)** Leave one out cross validation (LOOCV) for conditions where all ligands were present in at least two conditions (20 of 23) showed significant Pearson correlation between prediction and data. **d)** The correlation within conditions for LOOCV was consistently lower than the train fits but significantly outperformed models trained with scrambled condition labels, statistic calculated using two sided Mann–Whitney U test (n=20). **e)** The model was unable to fit some of the TFs and these also performed poorly under cross validation. **f)** Predicted effect on RELA activity in response to simulated KO under the TNFα stimulated condition using the LOOCV models to estimate consistency, top 10 KOs by average effect size, whiskers. Abbreviations: glucocorticoids (GC), ultrapure lipopolysaccharide (upLPS), standard LPS (LPSc), immune complexes (IC), Pam3CSK4 (P3C) and prostaglandin E2 (PGE2).

For the signaling network to accommodate the set of ligands used in the experimental study it was expanded with interactions from an immune system specific resource, InnateDB^38^ (see Methods). With this the model fit the data (Fig. 6b) with high accuracy. The data set was too small (23 samples x 58 TFs = 1334 datapoints compared to 7000> parameters) to be expected to generalize well, yet leave-one-out cross validation (LOOCV) showed that models trained on this data produced significantly better predictions than chance (Fig. 6c) or models trained on data in scrambled order (Fig. 6d). The generalization performance for these data was better than for synthetic data of comparable size, which may be due to denser sampling from a restricted region of the ligand stimulation space. We noticed IL4 among the conditions with poorer generalization, potentially due to confounding interactions with LPS^39^ that were not properly learned from the single condition available in the LOOCV setting. For some of the TFs the model did not produce acceptable fits (Fig. 6e), presumably due to limitations in the TF activity estimation or incompleteness of the signaling network.

In spite of these shortcomings, we simulated the effect of gene KOs on RelA under the TNFα stimulated condition (Fig. 6f), which is known to elicit long lasting effects on NF-κB signaling^35^. As could be expected, the TNF receptor was predicted to have the largest effect, but a number of NF-κB activating proteins were also identified, e.g. CHUK, IKBKB and RIPK1, in good agreement with prior knowledge, showcasing how biological relevance can be learned by this integrative approach. Still, the notable discrepancies between models suggests that much larger datasets are required.

## Discussion

We have demonstrated here that genome-scale simulation of intracellular signaling is now attainable. We developed a computational framework based on RNNs, constrained by prior knowledge of network interactions, to rapidly train predictive models of signaling using ligand-TF data pairs. In particular the model’s ability to predict the effects of KO’s is highly advantageous and cannot easily be matched by black-box based models. For models trained on real world data this would have important clinical implications, since many drugs act by blocking the activity of signaling molecules. We demonstrated the practical applicability of the framework on literature data and although this particular dataset contained too few conditions to generate reliable predictions, it showed how consistency between data and known biology can be enquired and highlighted some limitations with the prior knowledge networks. Our experiments with synthetic data suggest that the framework, in combination with larger datasets, could be used to train highly generalizable genome-scale models to further our understanding of intracellular signaling.

Presently, many high-throughput methods are being developed that will synergize with the framework, including large-scale transcriptomic screenings, e.g. the L1000^40^. These will enable profiling of numerous ligand-combinations to explore the space of possible signaling states and models trained on such data will provide a succinct and actionable representation of the acquired knowledge. The framework is not limited to study ligand-stimulation, our gene KO-simulations demonstrate how intracellular perturbations could be incorporated. Perturbations and KOs are of great interest for studying signaling^16,41^ and can help resolve identifiability issues, where multiple pathways fit the data equally well. Innovative use of nucleotide barcoding has enabled simultaneous construction of KO cells and sequencing of their gene expression^42^, albeit so far this was only applied to TFs, not signaling proteins. Identifiability issues could also be resolved by collecting data on the internal signaling states of select proteins, these data could be projected from the state vector and fitted analogously to TF data. High-throughput methods for generating multimodal data are currently under development, e.g. coupled profiling of transcriptome and protein activity^43^ and barcoding states of phosphoproteins using antibodies^44^. Alternatively, such data may be acquired using proteome level quantification of phosphorylation states^7^.

Transcriptomics is a strong technology for generating genome-scale data to train signaling models both in terms of cost and availability. Transcriptome based TF-activity estimates, provides a much larger set of observables than high level phenotype data, e.g. cell viability^16,19^, which is an alternative for large-scale signaling models. The connectivity of TFs throughout the signaling network also helps offset the increase in number of parameters with increasing network size by a matching increase in number of TFs, i.e. observed datapoints. The use of transcriptomics data to infer TF activity requires reliable estimation- methods. While many activities inferred using statistical methods are of high quality^11^, our understanding of gene regulation is continuously improving and more advanced computational methods are being developed, e.g. auto encoders that fit TF activities as latent variables informed by prior knowledge of TF-gene relations^45^ and mechanism-based deep learning models^34^. There is also development of sequencing based methods that simultaneously profile chromatin accessibility, intra-nuclear proteins, and gene expression^46^, which could aid in acquiring more accurate TF activity estimates. There are presently several methods that strive to infer signaling patterns from transcriptomics data and prior knowledge of the signaling- and regulatory networks, e.g. CARNIVAL^47^ and NicheNet^48^. However, these aim to provide qualitative descriptions of possible network wirings for individual condition as opposed to generating a predictive model consistent with all conditions as the one developed herein.

The framework relies strictly on prior knowledge of the signaling network and does not attempt to identify novel interactions. This is advantageous since it strongly reduces the solution space, alleviating the data requirements while at the same time enforcing biological plausibility. It also maps the hidden nodes to corresponding signaling molecules that allow KO events to be simulated. However, since it is not likely that the complete signaling network has already been characterized, the setup forces solutions to align with existing knowledge. At best this may result in inability to completely fit the test data, which may help highlight signaling interactions that require further attention and research, but incorrect relations may also be learned. One potential solution would be to allow the model to use a limited number of interactions supported by prior knowledge of sub-standard quality or that are completely novel, which could result in data driven discovery. This may be particularly relevant when expanding the framework to other mammals, since their prior knowledge networks often are mere homolog based extrapolations^9^ of the human signaling network. Contrarily, removing interactions from the prior knowledge network for non-expressed signaling molecules could aid the generation of cell type specific models.

We here aspire to model the effect of ligand stimulation in a single cell type and differentiation state. This is encoded in the constant weights and biases, but a natural generalization would be to let these parameters depend on external factors, e.g. cell type or test subject. Assuming that the wiring is mostly conserved, these parameters could be made into regularized functions of easily quantifiable properties, e.g. genotypes, allowing personalized parametrizations that still leverage data from other experiments. Subsets of parameters could even be pretrained using data from molecular studies, which corresponds to transfer learning that has been successful in other ANN applications, e.g. mammograms have been analyzed by appending a classifier to a network pretrained on regular images^49^. This would be particularly useful for analyzing subpopulation-specific responses among cells within a single experiment, that are now being inquired with single cell sequencing techniques. It is of much interest to discern the root cause of these differences e.g. differences in ligand concentrations, basal activity or network wiring. Single cell sequencing has founded an atlas of cell types at various stages of differentiation, and a fruitful continuation of this work, in particular for immune cells^50^, could involve comparisons of their dynamic responses to stimulation in through differences between parameterized models. Our framework relies on steady state assumptions motivated by time scale separation. From biological perspective it seems plausible that evolution would favor reproducible responses, i.e. that a given signaling pattern converges to the same state each time, although there are certainly exceptions, where sustained oscillations are instead desirable, e.g. the circadian rhythm or the cardiac cycle. Due to the steady state assumption the framework is not suitable to simulate such dynamics, although the RNN internally calculates a trajectory for each condition, these cannot be directly interpreted as time-series predictions. This is partly due to the initialization at zero, a biologically implausible state from which even the control condition, without ligand stimulation, is driven by basal activity from the bias term. But also, because no observations are provided along the trajectory before reaching steady state. For practical purposes this is not a concern, since the model predictions do not depend on the trajectories, but cellular dynamics may still be of interest for some applications. Time-series data could in principle be accommodated by the framework by fitting states at particular time steps, however, they would likely be better accommodated by a continuous time RNNs. Such RNNs have obvious similarities with ODE models and discussions on bridging the gap between RNNs and ODEs are ongoing^16^, notably a direct correspondence has been established between RNNs with a specific architecture and a common numerical ODE solver^51^.

Our regularization of the spectral radius ensures that all conditions in the training data reach steady state, but does not guarantee that this holds for arbitrary conditions. The pursuit of methods to enforce global stability for non-linear systems is an ongoing^52^, but it is not clear if global stability should be required for biological systems that may be unstable for conditions that are never encountered. Interestingly, if evolution is viewed as an optimization algorithm that has learned cellular parameters from conditions that are encountered, then by analogy turbulent states could be expected to occur for untrained conditions, which may be an interpretation of the chain-of-events in some diseases e.g. the detrimental immune responses known as cytokine storms^53^.

The challenge to learn parameters of a model with known structure from data is not limited to biology. In control theory, it has been proven that stochastic gradient decent (SGD) can learn linear dynamical systems^54^, which corresponds to an RNN with linear activation function. The RNN developed herein is an example of a sparse ANN. It has been recognized^21^ that for fully connected ANNs trained on image data, most parameters can be set to zero without marked loss in performance. After removing theses interactions, the sparse models can sometimes be retrained to the same level of performance as the original, since the learned structure remains encoded in the sparse connections. For the signaling network, sparsity has been learned by evolution. The ongoing development of new algorithms and hardware for training ANNs assures that the future will provide further improvements in model sizes, and training and execution times, e.g. sparse matrix multiplication is parallelizable and can be efficiently calculated on graphic processing units (GPU)^55^.

There are many avenues to expand the framework to further accommodate realistic simulations. One would be to allow molecules in different cellular compartment to have distinct signaling states. This would add a spatial component to the model and could be implemented directly through the prior knowledge network without changes to the framework. The intrinsic modularity of ANNs allow for intuitive integration with other networks, this seems immediately promising for integration with ANNs of regulatory processes, but it is also conceivable that cell-cell interactions could be modeled by chaining together multiple networks. The use of executable models in cancer research has shown how submodules with varying levels of abstraction can be integrated into a computer program that can be formally verified^56^. The rapid execution of trained models in consort with databases of drug-interaction partners^57^ opens up for genome-wide *in silico* screening of drug responses. This, together with personalized signaling models could provide individualized predictions of drug responses and side effects at the level of individual cell types.

## Methods

### Ordinary differential equations of molecular dynamics

ODEs were formulated for the different reaction schemas (See Fig. 1a) assuming mass action kinetics (See supplementary Fig. S1 for an example). The rate constants were manually parametrized (See Supplementary Table S1 for values) to yield sensible output. The differential equations were solved numerically using an initial value problem solver for systems of ODEs (scipy.integrate.solve_ivp^58^ in python 3.7.10). State variables were initialized as 1/[total number of states] and the activity after 100 time units was taken as the steady state value. For convenience the system was solved once with high resolution, a 50×50 linearly spaced grid, and linear gridded interpolation (scipy.interpolate.interpn) was used to down- sample to the indicated operational resolution.

### Neural network simulations of molecular interactions

Neural networks where constructed and trained using the pytorch framework^59^. This includes the autograd functionality, i.e. automatic differentiation, that retains the computation graph and uses it to automatically calculates gradients of the loss function. For the sigmoid activation the default formulation was used (torch.sigmoid), for ReLU the leaky version was used (torch.nn.functional.leaky_relu), and the MML function was manually implemented (as specified in Fig. 1b). For the fully connected layer (torch.nn.Linear) 5 hidden nodes were used. A trainable scaling factor was added to the output of the functions to accommodate normalization of activities. The neural networks were trained for 5000 epochs using the ADAM optimizer (torch.optim.Adam) with a learning rate of 0.002 and the built in L2 weight decay (factor 10^−5^). Default initialization of weights and biases was used.

### Structure of prior knowledge network files and ligand input and TF-output files

The signaling network structures were stored in list format with each entry containing a source node, a target node, the mode of action, and references to databases and PubMed ids, where applicable. Signaling nodes were identified by their uniprot identifier. This structure is similar to the format used by OmniPath^9^, but unlike OmniPath, all interactions were considered directed from source to target and reversible interactions were represented by an additional entry with source and target nodes exchanged. The signaling network file was accompanied by an annotation file, that for each of the signaling nodes specified their function, e.g. ligand or transcription factor, and a human readable synonym, e.g. gene name or small molecule acronym. For storage of trained networks pytorch serialized objects (torch.save) were used and a human readable plain text format was also developed where each entry contained the parameter type (bias, weight, input projection or output projection), parameter value, source node and target node (only used for weights). For the macrophage dataset input and output data for the network were stored as tab separated tables with conditions as rows and ligands and TF levels respectively as columns.

### Projections of matrices from input to state and from state to output

Input consists of a [*s* × *i*] matrix where *s* is the number of samples (in total or in the mini-batch) and *i* is the number of ligands in the input, the output consists of a [*s* × *o*] matrix, where *o* is the number of TFs in the output. The recurrent neural network calculates a state matrix, [*s* × *n*], where n is the number of state variables. To accommodate size differences between input, output and state matrixes the RNN is proceeded by a projection layer that inserts the elements of the input at their corresponding position in a zero-padded matrix [*s* × *n*] with elements ordered as in the state matrix. Similarly, the state vector is projected to an output matrix by selecting the corresponding TF elements from the state matrix and placing them in an order that matches the order of TFs in the data. Scaling factors for each element are included in the projections and for the output projection these are made trainable parameters.

### Recurrent neural network formulation

The recurrent neural network takes a matrix *b*_*in*_ as input and returns a matrix *x*_*ss*_ as output both with the structure [*s* × *n*], with *s* and *n* defined as above. The function is parameterized by trainable weight and bias vectors. The structure of the signaling network (*A*) is provided as a sparse row matrix (scipy.sparse.csr_matrix) with values of the non-zero elements given by the weight vector. The columns of the matrix correspond to sources and the rows to targets. The state vector is initialized as all 0 and iterated for a finite number of steps, set to 150 in this study. The RNN function was implanted as a manual autograd function (torch.autograd.Function) with both forward and backward pass specified manually (see Algorithm 1 in Supplementary methods) using numpy^60^ operations. The spectral radius of the transition matrix for the backward pass is assumed to be less than 1, meaning that the magnitudes of the back propagated gradients are bounded. However, since it cannot be excluded that this constraint occasionally will be violated during training, gradient clipping is applied at each iteration. To prevent clipping under regular conditions, the clipping function was constructed with a linear segment between two saturating tanh regions (See supplementary materials algorithm 1).

### Initialization of weights, biases and scaling factors

Weights are initialized as *u*(0, 0.1) + 0.1, where *u* is a uniformly sampled random number on the specified interval. Weights corresponding to inhibitory interactions are made negative by multiplication by −1. All weights are scaled by a factor 0.8/*ρ*(*A*), where *ρ*(*A*) is the spectral radius of the matrix A to ensure that *ρ*(*A*) < 1. Biases are initialized at a value of 0.001 except for biases corresponding to nodes that only have inhibitory inputs, in which case they are initialized at 1 to accommodate dynamic node states in the positive regime. The scaling factors for elements in input and output projections are initialized by the same value, 3 for input projections (which corresponds to a state of ∼0.92 after applying the activation function) and 1.2 for output projection.

### Soft constraints for weight sign and ligand bias

To impose soft constraints, barrier functions were constructed, multiplied by a constant and added to the loss function. For interactions with known mode of action, activation or inhibition, the sign of the corresponding weight was constrained to be positive or negative respectively. This was imposed by adding the sum of absolute values of weights where the sign conflicted with prior knowledge. For biases associated with ligands, the model was prevented from learning large values, since knowledge about ligand concentrations is expected to be available and provided as input. Here, the barrier function was constructed as the sum of squares of biases belonging to ligands.

### State and parameter regularization and application of state noise

To aid in generalization and prevent the model from getting stuck in local minima, several regularization techniques were applied. To prevent parameters from taking on extreme values, L2 regularization of weight and bias parameters was implemented by adding the sum of squares of these vectors multiplied by a coefficient, 10^−8^, to the loss function. For training on the synthetic dataset and the macrophage data an additional term was added to the weight loss to prevent weights from getting stuck at zero,

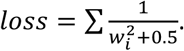

Regularization of the state variables was implemented to ensure that they remained active with a wide dynamic range during training, with similar objective as batch normalization. The goal was for each of the states to have a uniform distribution across conditions, and this was implemented by regularizing some of the statistical properties to match the corresponding properties of a uniform distribution on the interval [a b], i.e.

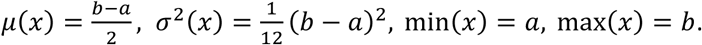

To be operational independent of batch size, the properties were calculated across all conditions, and for conditions that were not in the present batch their latest calculated values were used, however these were detached (torch.tensor.detach) from the computation graph and only gradients from the current batch were back propagated. The regularization was implemented by calculating the deviation of the empirical property across conditions from the ideal property calculated for the interval [0.01, 0.99]. The sum of squares of deviations was applied as barrier function. Additional regularization was added to prevent negative max values, the sum of negative max values was used as barrier function. The sum of these contributions was multiplied by a factor, 10^−5^, and added to the loss function.

In addition to regularization, gaussian noise was added to the *b* vector for each forward pass, to ensure that the fitted parameters were robust to small deviations. The level of noise was made proportional to the learning rate (lr) as *b* = *b* + 10 · *lr* · *norm*(0,1), where norm is sampling from a normal distribution with 0 mean and 1 variance.

### Spectral radius regularization

An exponential barrier function was used to constrain the spectral radius (*ρ*)

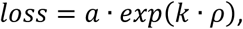

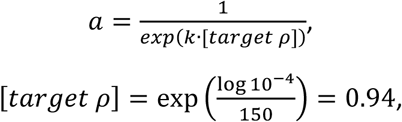

with *k* = 21 and a scaling factor *a*, where 10^−4^ is the target precision and 150 the number of steps in the RNN. To be able to backpropagate this function, we constructed a manual autograd function for the gradient of the spectral radius (see supplemental methods Algorithm 2). It made use of made use of a sparse eigenvalue solver (scipy.sparse.linalg.eigs). Since both left and right eigenvectors are required for the gradient calculation and only right eigen vectors are returned by the sparse solver, the matrix was transposed and solved a second time with the predicted eigenvalue from the first pass as target. To conserve computations, a single steady state was selected at random for regularization from each batch.

### Model reconstruction

The most recent interaction database was retrieved from the OmniPath^9^ website (archive.omnipathdb.org, retrieved 2021-06-21). Only human interactions from the OmniPath core set were included. The interactions were divided into 3 subsets, Ligand-Receptor (LR) interactions, regulatory interactions and signaling interactions (see supplementary table S3 for details on the queries). The LR and signaling interaction were further reduced to only include interactions that referenced KEGG among the sources. A few reactions were removed based on manual curation; interactions between IL6R and JAK1, STAT3 and SRC were removed, since IL6R only signals through its interaction with gp130 (IL6ST)^61^; the interaction between TLR4 and IRAK4 and CD14 were removed from receptor-ligand interactions, since IRAK4 and CD14 are not considered ligands based on their uniprot annotation^62^. All interactions that were listed as reversible were duplicated and reversed and their interactions were set as unidirectional. To avoid duplicate interaction, all interactions present in LR were removed from the signaling set. Any conflicts in mode of action, i.e. listed as both activating and inhibitory, were resolved by removing the mode of action information. And nodes without mode of action information were removed. Nodes that were not listed in Uniprot^62^ were also removed. Nodes were classified as ligands if they were listed as sources in LR, as receptors if they were listed as targets in LR and TFs if they were listed as sources in the regulatory interactions. The LR and signaling interactions were merged. Nodes where considered dead ends and removed from the network if there for the node was no path from any ligand or to any TF. Additionally, nodes were considered redundant and removed if they had only a single source and target that both were the same node. Network plot was drawn using MATLAB 2020a.

The same procedure was followed for the network intended for the experimental data, but InnateDB was also included as an approved source. Furthermore, the RL interactions were manually defined (supplementary table S3) based on the ligands available in the experimental data based on uniprot^62^ annotation. The list of TFs was restricted to the ones with experimental data available.

### Synthetic data generation and analysis

To fit parameters that result in non-trivial and biologically plausible predictions, an objective function was defined including several terms. For each epoch the model was provided with 200 conditions containing 5 randomly selected ligands per condition with uniformly sampled concentrations, these were resampled for each evaluation. The predicted TF activities were regularized to follow a uniform distribution both across conditions and across different TFs, this was implemented in the same way as state regularization (See above) but without dependency on states from previous epochs. The mean correlations were minimized across both conditions and TFs, this was implemented by calculating the average of the correlation coefficients of the output matrix and its transpose. Spectral radius regularization was applied with a coefficient of 10^−2^ and L2 norm on weights and biases was applied with a coefficient of 10^−6^. Sign and ligand constraints were applied as specified above. To preempt information leakage, parameters were initialized differently than for the trained models; weights were initialized uniformly at random from the interval [0, 3] and their sign was assigned based on mode of action, and scaled to a spectral radius of 0.8; bias was assigned by sampling uniformly from the interval [0, 0.01].

The complexity of the synthetic data was studied by PCA analysis of predicted output from 2000 randomly generated conditions with different numbers of simultaneous ligands. Linear models (sklearn.linear_model.LinearRegression)^63^ were fitted to the synthetic data for each simultaneous ligand level, and prediction performance was evaluated using 10 fold cross validation (sklearn.model_selection.KFold). The performance of the model trained on single ligand data was also evaluated.

### *In silico* knock outs

The predicted effects of *in silico* KO were studied by adding a strong negative bias (−5) to the node of interest, resulting in near zero node states after applying the activation function. The change in TF activity compared to the control condition without KO was used as metric since most TFs are not expected to be affected by most KOs. For the KO predictions under the TNF condition, the KO was applied to all nodes in each of the models that were generated for the LOOCV, and nodes were ranked by the average of the predicted effect on RelA.

### Inference of TF activity

Literature data^4^ was retrieved from the ArrayExpress^64^ database (ebi.ac.uk/arrayexpress), accession number E-GEOD-46903. Genes without any detected signal (min p>0.01) or without variance ([std] < 10^−6^ [mean]) were removed from further analysis. The log-transformed data (5203 genes and 384 samples) was centered and TF activities were inferred using Dorothea^11^, an enrichment based statistical method. Only TFs with a confidence score of A or B and interacting with at least 5 genes were included. Conditions were filtered to only contain data from GM-CSF cultured macrophages from the same time point (72 h) amounting to 103 samples including biological replicates. The Dorothea reported log odds ratios were transformed to probabilities using the inverse logit function, i.e. the logistic function. The average was taken among replicates resulting in 23 unique conditions. Standard deviation among replicates for each TF within each condition were inspected and TFs were discarded if their 75^th^ percentile of standard deviations exceeded 0.2 corresponding to 4 TFs (See supplemental Fig. S7a).

### Hardware for simulations

Simulations were performed on a Dell Precision 3530 laptop with an Intel i7 CPU @ 2.60 GHz with 6 cores (12 logic processors) and 16 GB ram. For convenience, evaluation of data requirement and cross-validation was carried out on a singled threaded computer cluster (Intel Xeon CPU @ 2.60GHz) that allowed job scheduling (using Slurm) with 16 parallel jobs.

## Data and code availability

The code and scripts the to reproduce the simulated data are made available through a public repository (github.com/Lauffenburger-Lab/Artificial-Signaling-Network).

## Acknowledgments

The authors want to thank Eduardo Sontag, Shu Wang, Filip Buric, Carolin Loos, Chuangqi Wang, Brian Joughin, Lauren Baugh, Dan M Schafer, Adityanarayanan Radhakrishnan, Caroline Uhler and Karen Sachs for valuable input. We acknowledge funding from Vetenskapsrådet (AN, 2019-06349) and National Institutes of Health (NIH) (DAL, AI-201700104; JMP and BB, R01A1022553, R01AR073252 and BAA-NIAID- NIHAI201700104).

## Author contributions

AN and DAL conceived the study and JMP, BB and provided input on the study design, AN implemented the code and executed the simulations, AN wrote the manuscript, JMP, BB and DAL edited the manuscript.

## Competing interests

The authors declare no competing interest.

## Notes

https://github.com/Lauffenburger-Lab/Artificial-Signaling-Network

### Competing Interest Statement

The authors have declared no competing interest.

