## Supplemental materials for "Artificial neural networks enable genome-scale simulations of intracellular signaling"

### Supplementary Figures

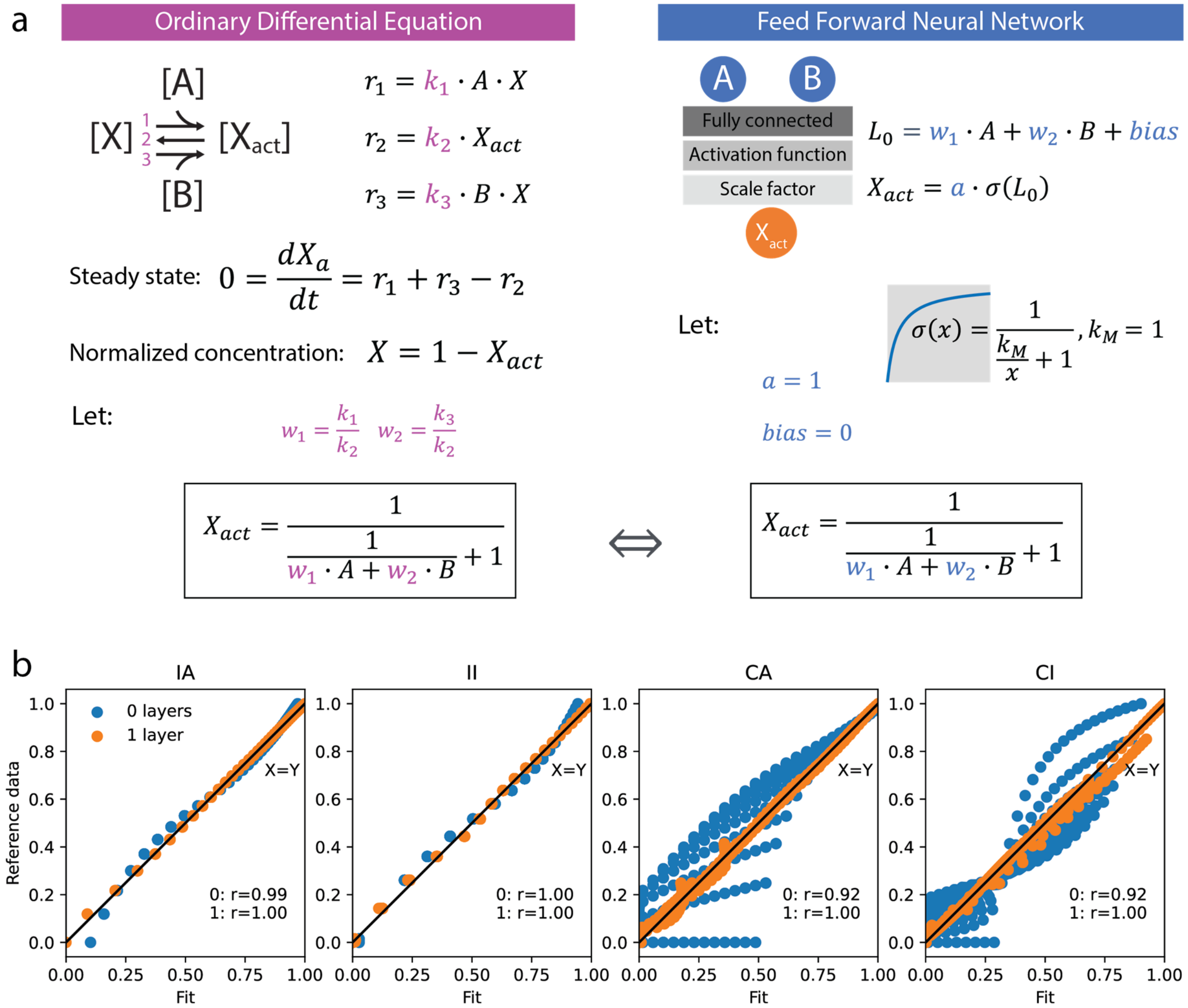


**Supplementary Figure S1 Fitting ODEs with a Michaelis Menten like (MML) activation function. a)** The analytical solution of steady state activity for independent activation derived from the ordinary differential equations. The equation is equivalent to the equation for a zero-layer neural network with the Michaelis Menten equation as activation function. **b)** Neural network approximations of ODEs with different mechanism using an MML activation function, IA=independent activation, II=independent inactivation, CA=cooperative activation, CI=competitive inhibition.

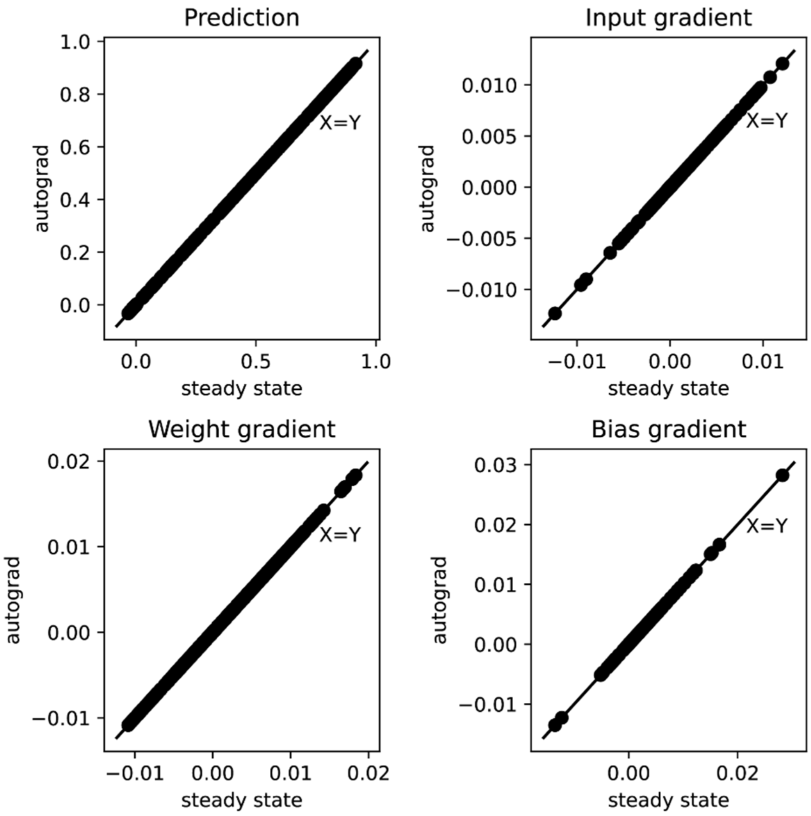


**Supplementary Figure S2 Numerical comparison of auto-grad and steady-state for a random network.** The predictions and the gradients for input, weights and bias are indistinguishable between models computed by pytorch autograd (automatic differentiation) function and the manually implemented steady-state function. This function only depends on the steady state gradients and thus has significantly lower memory overhead compared to autograd that stores the full computation graph in memory and consequently, autograd takes approximately 10 times longer to complete. This also means that arbitrarily many timesteps can be executed without running in to memory issues.

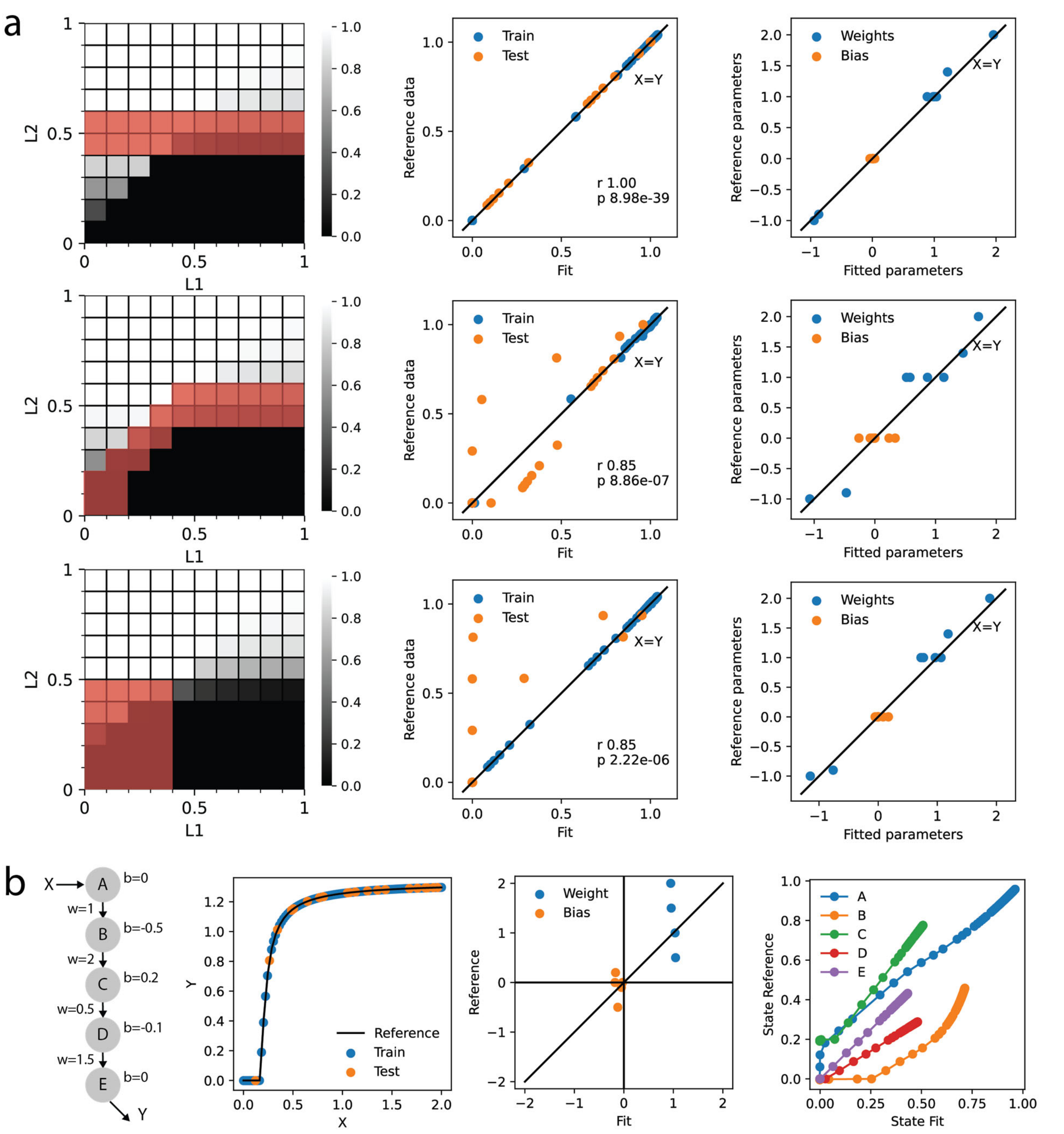


**Supplementary Figure S3 Adversarial test data and network structure. a)** The networks ability to extrapolate was test by purposefully selecting training data in challenging locations, including along the boundary and in the whole bottom left quadrant. **b)** An unbranched network structure was constructed and a reference model was manually parameterized. The test fit to this model was perfect, but parameters were not accurately predicted. The node states of the fit and reference were correlated.

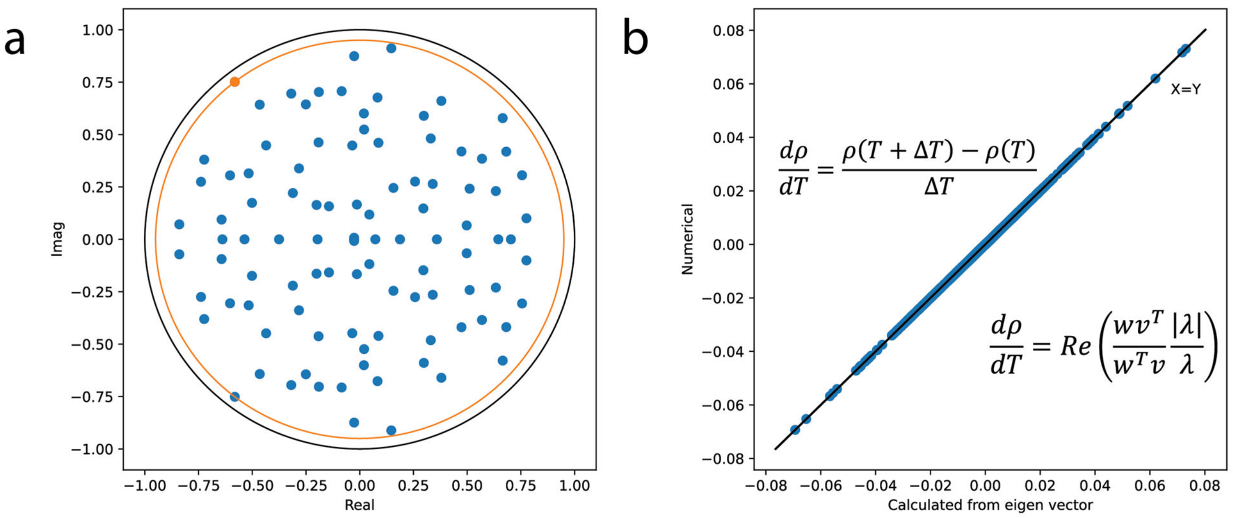


**Supplementary Figure S4 Derivative of spectral radius. a)** Eigen values for a randomly parameterized transition matrix, where the largest absolute value of the eigen values (orange dot) is the spectral radius. Note that due to the transition matrix being real and non-symmetrical eigenvalues are mirrored along the origin of imaginary axis. **b)** Comparison of the derivative of the spectral radius calculated from its left and right eigen vectors and numerically calculated by perturbing each element of the transition matrix in turn by a small value (epsilon= $10^{-10}$) and re-calculating the eigen values.

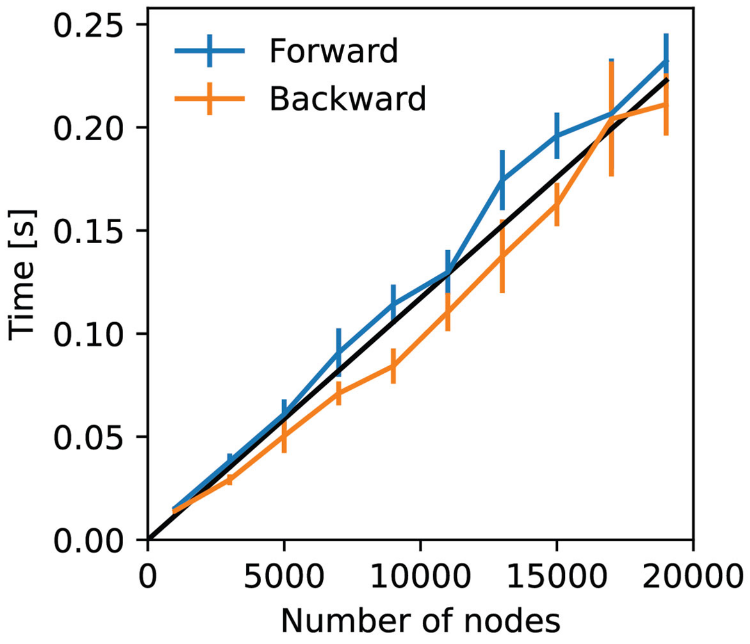


**Supplementary Figure S5 Test of wall time for different network sizes.** Sparse random networks with 10 interactions per node were generated (using scipy.sparse.random) and their spectral radius was constrained to 0.9. The average wall time (of 5 random networks) for the forward and backward pass was calculated using random input and output (with batch size of 3). Error bars show standard deviation. There was a linear fit (R2=0.96) between number of nodes and time.

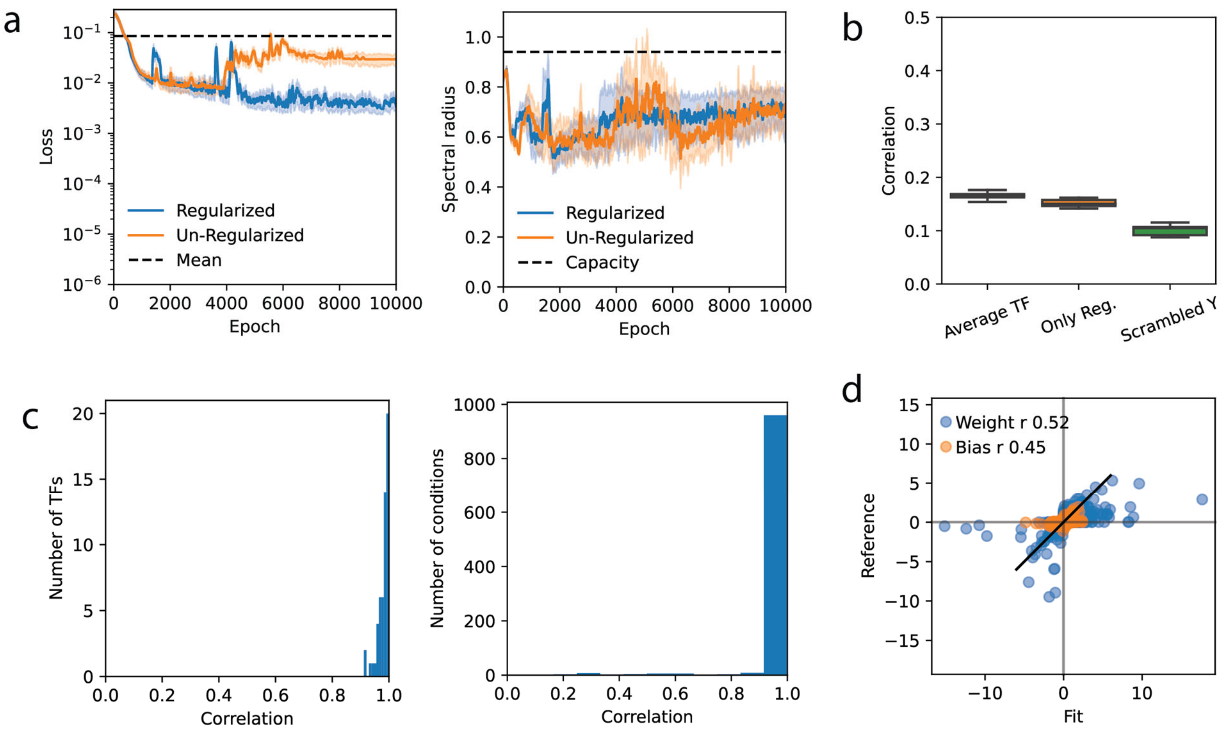


**Supplementary Figure S6 Training on a synthetic dataset. a)** Training trajectories for two models, one with and one without spectral radius regularization trained on data from the same 10 conditions. Without spectral regularization training diverges (around epoch 4000) and the spectral radius sometimes increases above capacity, which is a value (less than 1) that depends on the number of timesteps. **b)** Comparison of fits of a prediction consisting of the average of each TF, a model trained using only the regularization (Reg.) terms, and a model trained with scrambled condition order. 10 tests consisting of 1000 conditions each were sampled at random. **c)** Correlation between model and reference for individual TF and individual conditions evaluated on 1000 test conditions. **d)** Comparison between reference and fitted parameters, note that the correlation for weights is heavily influenced by the sign that is constrained by regularization. The correlation for the absolute value that is not influenced is 0.36, p $10^{-27}$.

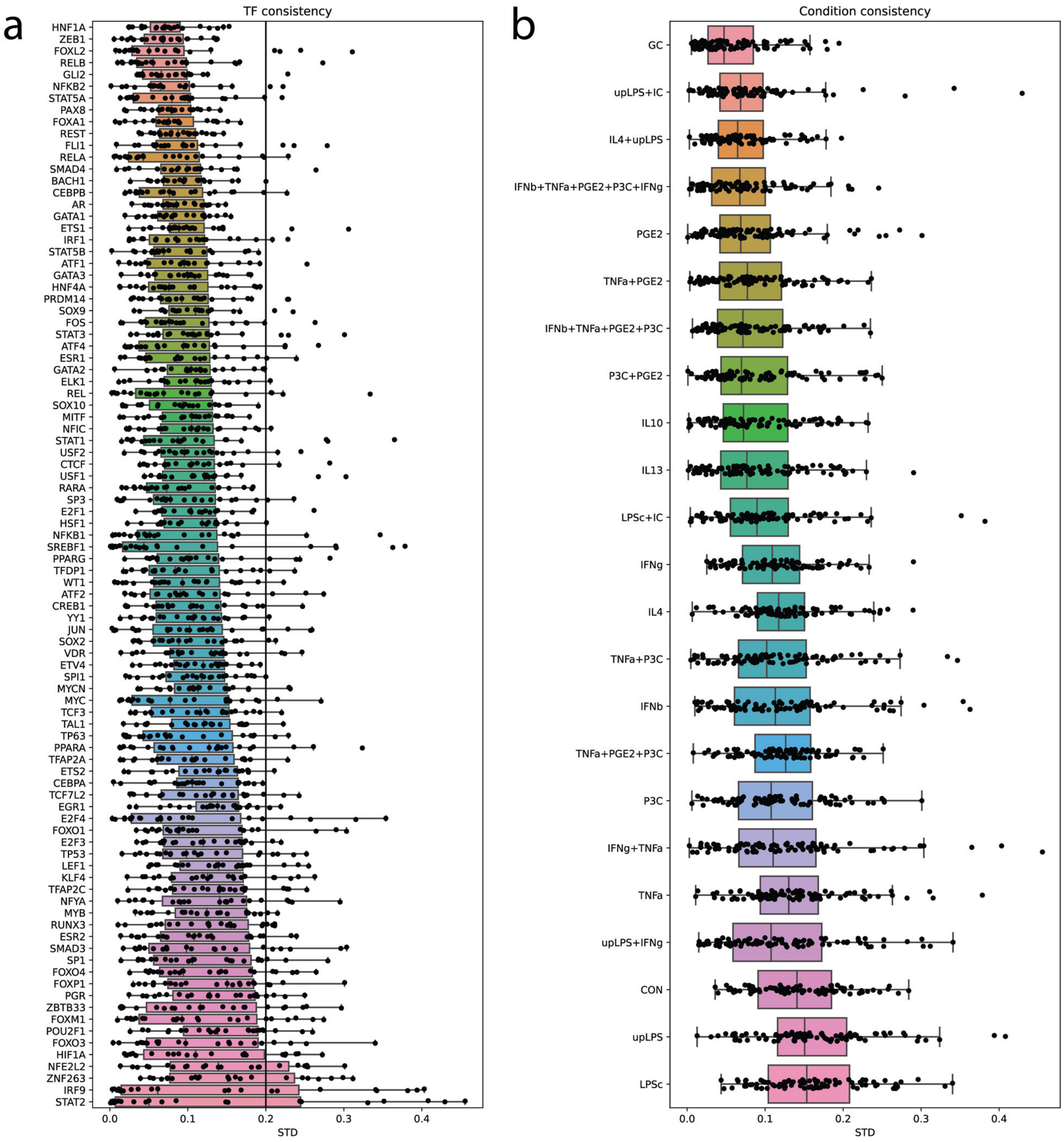


**Supplementary Figure S7 TF activities inferred from experimental data. a)** Standard deviation of TF activities among replicates by transcription factors across conditions. A cutoff value of 0.2 (black line) for the 75th percentile was selected after inspecting the graph. Four transcription factors did not meet the threshold, STAT2, IRF9, ZNF263 and NFE2L2. **b)** As A, but by condition across TFs.

### Supplementary Tables

**Supplementary Table S1 Rate parameters for different molecular mechanisms.**

| | $k_1$ | $k_2$ | $k_3$ | $k_4$ |
|---|---|---|---|---|
| **IA** | 0.2 | 0.1 | 0.2 | NA |
| **II** | 0.2 | 0.1 | 0.2 | NA |
| **CA** | 0.5 | 0.1 | NA | NA |
| **CI** | 0.5 | 0.1 | 2 | 0.1 |

NA = not applicable

**Supplementary Table S2 Queries used to prune the interactions in the OmniPath database**

| Purpose | Field | Operation | Value |
|---|---|---|---|
| Only human | ncbi_tax_id_target | == | 9606 |
| Only OmniPath core set | omnipath | == | True |
| Ligand-Receptor (LR) interactions | Ligrecextra **AND**<br>sources | ==<br>contains | True<br>'KEGG' |
| Signaling interactions | post_translational **AND**<br>sources | ==<br>contains | True<br>'KEGG' |
| Regulatory interactions | dorothea **AND**<br>dorothea_level | ==<br>contains | True<br>'A' **OR** 'B' |

**Supplementary Table S3, manually defined RL interactions based on uniprot annotation.**

| Source | Target | Motivation |
|---|---|---|
| GC | NR3C1 | Glucocorticoid receptor |
| IC | FCGR3A | Immune complexes may activate FC receptors |
| IC | FCGR2A | |
| IFNb | IFNAR1 | Interferon alpha/beta receptor 1&2 |
| IFNb | IFNAR2 | |
| IFNg | IFNGR1 | Interferon gamma receptor 1&2 |
| IFNg | IFNGR2 | |
| IL4 | IL4R | Interleukin-4 receptor |
| IL10 | IL10RA | Interleukin-10 receptor subunit alpha |
| IL13 | IL13RA1 | Interleukin-13 receptor subunit alpha |
| P3C | TLR2 | P3C activates TLR1 & 2 |
| PGE2 | PTGER3 | PGE2 activates PTGER1-4 |
| LPSc | TLR2 | LPS activates cells through TLR4, the TLR2 activity of LPS-PG is ascribed to a contaminant lipoprotein |
| LPSc | TLR4 | |
| LPSc | CD14 | CD14 binds bacterial lipopolysaccharide |
| upLPS | CD14 | |
| upLPS | TLR4 | Ultrapure LPS only activates TLR4 |
| TNFa | TNFRSF1A | Tumor necrosis factor activates receptor 1 A and B |
| TNFa | TNFRSF1B | |

### Supplementary Methods

#### Algorithms

The algorithms presented below (Algorithm 1 and 2) are manual autograd functions and thus require both forward calls and backward calls (partial derivative). To accommodate dot multiplication, data must be transposed to [features x samples], as compared to the machine learning standard of [samples x features]. In practice this is performed by the algorithm but for the sake of this presentation it is assumed to already have occurred.

---

**Algorithm 1 (forward): Sparse one-to-one RNN**

[$x_{ss}$, $x_{raw}$] = **function** forward($b_{in}$ weights, bias, A)

**Input**:

$b_{in}$, [n x s] matrix for s samples and n state variables, zero-padded with ligand concentrations at their corresponding positions.

weights, [i x 1] vector containing the non-zero elements.

bias, [n x 1] vector of biases for each state variable.

A, [n x n] sparse matrix with i non-zero elements that describes the network structure.

**Output**:

$x_{ss}$ [n x s] vector of predicted steady states.

$x_{raw}$ [n x s] vector of predicted steady states before applying activation function.

---

A.nze = weights *#loads the weights into their correct positions in the sparse matrix A*

b = $b_{in}$ + bias

$x_{ss}$ = zeros [n x s] #initiate $x_{ss}$ as an all zero matrix with dimensions n x s

**for** max_iter **steps**: *#Max_iter is set to 150 in this study*

$x_{ss}$= **dot product**(A, $x_{ss}$)

$x_{ss}$= $x_{ss}$**+** b *#vector b is repeated for each sample in $x_{ss}$(broadcasted)*

$x_{ss}$= **activation**($x_{ss}$) *#element-wise activation function, specified above*

$x_{raw}$ **= dot product**(A, $x_{ss}$) **+** b *#When $x_{ss}$ is at steady state, this is same as inv(activation($x_{ss}$))*

**Algorithm 1 (backward): Sparse one-to-one RNN**

[$z$, dw, db] **= function** backward($err$, $x_{ss}$, $x_{raw}$, A)

**Input**:

$err$, [n x d] matrix containing gradient back propagated from comparing TF activities projected from $x_{ss}$ to data.

$x_{ss}$, [n x d] matrix see forward.

$x_{raw}$, [n x s] matrix see forward.

$A$, [n x n] matrix see forward.

**Output**:

$z$, [n x s] error to be back propagated to proceeding layers.

dw, [i x 1] gradient of the weight vector.

db, [n x 1] gradient of the bias vector.

*xDelta* **= activation'(**$x_{raw}$**)** *#the derivative of the activation function*

T = **transpose**(A) #targets and sources are reversed for backward pass

z = An all zero matrix with same dimensions as $err$

**for** max_iter steps:

z = **dot product**(T, z) + err

z = **element-wise product**(xDelta, z)

z = **clipping**(z) *#gradient clipping as described below*

db = **rowsum**(z)

dw = **rowsum**(**element-wise product**($x_{ss}$[source], z[target])) *# as further explained below*

For calculation of the weight gradients, only the non-zero elements of A are relevant. Therefore, the complete dot multiplication between the matrixes $x_{ss}$ and grad can be replaced by elementwise multiplication and subsequent summation for the elements corresponding to the source (column) and target (row) of each weight.

To prevent clipping under normal conditions, the clipping function is constructed with a linear segment between two saturating tanh regions, resulting in a continuous and monotonically increasing function from -2 to 2,

$$
\begin{array}{cc}
 & clipping(z) \\
z \leq -1 & \tanh(z+1)-1 \\
-1 < z \leq 1 & z \\
1 < z & \tanh(z-1)+1
\end{array}.
$$

**Algorithm 2 (forward): Spectral radius**

[$\rho$, $e$, $v$] = **function** forward(A, $\overline{weights}$)

**Input**:

A, [n x n] sparse matrix with i non-zero elements that describes the network structure.

$\overline{weights}$, [i x 1] vector of linearized non-zero weights of A

**Output**:

$\rho$, scalar, the spectral radius

$e$, complex scalar, the eigen value of A with largest absolute value

$v$, [n x 1] complex vector, the right eigen vector of A

```
T = A                   #Create T with same structure as A
T.nze = weights         #load the linearized weights into correct positions in the sparse matrix T
e, v = eigs(T)          #returns the eigenvalue (e) with largest absolute value and its right eigenvector (v)
ρ=abs(e)
```

**Algorithm 2 (backward): Spectral radius**

[dw] **= function** backward(grad, $v$, $e$)

**Input**:

grad, a scalar with the backpropagated gradient of the barrier function

$v$, [n x 1] complex vector, see forward

$e$, complex scalar, see forward

**Output**:

dw gradient of the weight vector

```
e, w = eigs(tanspose(T), e) #gives the eigen vector w closest to the eigenvalue to e (the left eigen vector)
divisor = dot product(transpose(w), v)
direction = e/abs(e)
delta = elementwise product (w[target], v[source])/divisor
delta = real(delta/direction)
delta = limit norm(delta, 10) # optionally the norm of delta can be constrained to a finite value
dw = grad * delta
```

Since eigs is a stochastic algorithm it sometimes, under rare circumstances, fails to return eigen values before timing out, then zeros were returned as gradient.

#### Derivation, backpropagation at steady state

Let the state at time step n be defined as,

$$z_n = Ax_{n-1} + b$$

$$x_n = \sigma(z_n)$$

$$y_n = px_n.$$

Apply the chain rule to some loss function L($y_n$),

$$\frac{dL}{z_n} = \frac{dL}{dx_n}\frac{dx_n}{dz_n} = \frac{dL}{dx_n}\,\sigma'(z_n).$$

where

$$\frac{dL}{dx_n} = \frac{dL}{dy_n}\frac{dy_n}{dx_n} + \frac{dL}{dz_{n+1}}\frac{dz_{n+1}}{dx_n} = \frac{dL}{dy_n}\frac{dy_n}{dx_n} + \frac{dL}{dz_{n+1}}A$$

Combining the two equations we get the following recursive formula

$$\frac{dL}{dz_n} = \left(\frac{dL}{dy_n}\frac{dy_n}{dx_n} + \frac{dL}{dz_{n+1}}A\right)\,\sigma'(z_n).$$

If we assume steady state,

$$z_{ss} = Ax_{ss} + b$$

we get the following linear equation system

$$\frac{dL}{dz_{ss}} = \left(\frac{dL}{dy_{ss}}\frac{dy_{ss}}{dx_{ss}} + \frac{dL}{dz_{ss}}A\right)\,\sigma'(z_{ss})$$

expressed in gradient form

$$\nabla z_{ss} = \sigma'(z_{ss}) \odot (A^T \nabla z_{ss} + \nabla L\ ).$$

This equation can be solved by iteration from some initial guess for $\nabla z_n$ (e.g. $\nabla z_n = 0$), or using a numerical solver. From this the gradients of the parameters A and b can be calculated with respect to L as

$$\frac{dL}{dA} = \frac{dL}{dz_{ss}}\frac{dz_{ss}}{dA} = x_{ss}\nabla z_{ss}^T.$$

$$\frac{dL}{db} = \frac{dL}{dz_{ss}}\frac{dz_{ss}}{db} = \mathbf{1} \cdot \nabla z_{ss}.$$

If we prefer to not assume steady state at this stage, we can note that the loss function is evaluated at a specific timestep (k) and is zero elsewhere. The loss depends on states from timestep 1 to k and the loss is zero with respect to timesteps outside of this range.

$$\frac{dL}{dy_k}\frac{dy_k}{dx_k} = \nabla L, \frac{dL}{dz_0} = 0, \frac{dL}{dz_{k+1}} = 0, \nabla z_k = \sigma'(z_k) \odot (\nabla L)$$

With these boundary conditions the loss can be evaluated as

$$\nabla z_n = \sigma'(z_n) \odot (A^T \nabla z_{n+1}) = \left( \prod_n^k \sigma'(z_n) \odot A^T \right) \nabla L$$

Since parameters are the same across all time steps, their gradients can be summed into a quantity that is independent of the time step:

$$\frac{dL}{dA} = \sum_1^k \nabla z_i \, x_i = \sum_1^k \left( \prod_i^k \sigma'(z_i) \odot A^T \right) \nabla L x_i$$

$$\frac{dL}{db} = \sum_1^k \nabla z_i \cdot \mathbf{1} = \sum_1^k \left( \prod_i^k \sigma'(z_i) \odot A^T \right) \nabla L \cdot \mathbf{1}.$$

Note that the conserved part of these equations can be written in recursive form as

$$s_n = \sigma'(z_{k+1-n}) \odot A^T (s_{n-1} + \nabla L).$$

For large n the repeated multiplication will cause later terms to vanish, and the recursion can be truncated at the step (t) where this occurs. If forward propagation has reached steady state all of the non-truncated values will be from the steady state (and if not, then a larger k can be chosen, see next section for further analysis on the requirements for steady state to occur). We can then simplify the equation

$$s_t = \sigma'(z_{ss}) \odot A^T (s_t + \nabla L).$$

This has the same form as the steady state expression derived above, and the same analysis for parameter gradients applies.

### Derivation, spectral radius determines rate of convergence

Let the state at time step n be defined as,

$$x_n = \sigma(Ax_{n-1} + b)$$

then a steady state $x_{ss}$ is defined as

$$x_{ss} = \sigma(Ax_{ss} + b)$$

Taylor expand to the first order around the steady state

$$x_n = \sigma(Ax_{ss} + b) + \big(I\sigma'(Ax_{ss} + b)\big) \odot \big((Ax_{n-1} + b) - (Ax_{ss} + b)\big)$$
$$= x_{ss} + \big(I\sigma'(Ax_{ss} + b)\big) \odot \big(A(x_{n-1} - x_{ss})\big),$$

where $\odot$ is element wise multiplication and I the identity matrix. Let a displacement from the steady state at time step n be defined as

$$\Delta x_n = x_n - x_{ss},$$

and let $T$ be defined as

$$\big(I\sigma'(Ax_{ss} + b)\big) \odot A = T,$$

we can then rewrite the equation as

$$\Delta x_n = T\Delta x_{n-1}.$$

By repeated insertion we have

$$\Delta x_n = T^n \Delta x_0.$$

For an initial non-zero displacement from steady state $\Delta x_0$ to reach $x_{ss}$ at time n,

$$0 \approx T^n(\Delta x_0),$$

this requires that

$$T^n \approx 0,\ \rho(\mathrm{T}) < 1,$$

where $\rho(\mathrm{T})$ is the eigenvalue of T with largest absolute value. An approximation of the number of time steps (n) required depends on the desired precision and is given as

$$n \propto \frac{\log([precision])}{\log(\rho(\mathrm{T}))}.$$

To impose the spectral radius as a soft constraint, several forms for the regularization function were considered, including a reciprocal and exponential dependence on the spectral radius. A reciprocal function, while perhaps the most intuitive, undergoes a singularity at a spectral radius of 1 with infinite derivative, which is numerically impractical for gradient decent. An exponential function with parameters (a and k) could be fitted to behave similarly to the reciprocal function, but without singularity issues.

$$L_\rho = a\big(e^{(k\rho)} - 1\big)$$

### Analysis of the framework's algorithmic complexity

The computation of state vectors relies on sparse matrix multiplications that increases linearly with the number of interactions (z, number of elements in matrix) and squared by the number of signaling nodes (n, width/height of matrix). Because the number of interactions is expected to increase as a function of the number of nodes, the worst-case time complexity becomes $n^3$, corresponding to dense matrix multiplication. The calculation of spectral radius and eigen vectors of sparse matrixes used for regularization is naively of complexity $n^3$, but is calculated using a stochastic algorithm with a complexity $n^2$ and a factor that depends on the size gap[59] between the eigen values (g). This eigengap can be expected to be a function of n since the total number of eigen values (n) are constrained to a disk with finite area ($1^2\pi = \pi$). The model is iteratively evaluated until steady state, and computation time increases linearly with the number of time steps (t). Through matrix operations, different conditions can be evaluated in parallel, but the algorithmic complexity depends linearly on the number of conditions, that are typically divided into mini-batches (b) of some size (r). To train a generalizable model, the forward pass must be executed multiple times and each forward pass is accompanied by a backward pass with the same complexity. The number of epochs (e) required for the training to converge may depend on n and the number of conditions may in practice also depend weakly on n, as more parameters could be expected to require more data. With this notation the overall complexity depends on many factors,

$$O\big(er(btzn^2 + gn^2)\big),$$

an expression with explicit squared complexity with respect to n. However, due to relations between n and some of the other parameters, the complexity can be expected to be worse in practice.
